# Contingencies in biofilm adaptation of *Mycobacterium tuberculosis*

**DOI:** 10.1101/2024.06.07.597933

**Authors:** John F. Kernien, Madison A. Youngblom, Tracy M. Smith, Sydney S. Fry, Mohamed A. Mohamed, Holly J. Murray, Kadee N. Lawrence, Caitlin S. Pepperell

## Abstract

Biofilms are structured microbial communities that offer protection from a range of environmental stressors. Studies of experimental evolution within biofilms have yielded important insights into mechanisms of biofilm formation, as well as fundamental principles governing bacterial adaptation within these structured communities. Building on research using tractable species, our lab created a model for studying biofilm adaptation in *Mycobacterium tuberculosis (M. tb),* a fastidious, slow growing and lethal pathogen of humans. Here, we evolved eighteen *M. tb* populations arising from six parental genetic backgrounds under biofilm selection to investigate diversity in mechanisms of *M. tb* biofilm adaptation. We found that fine-scale differences among strains at the initiation of the experiment influenced the degree of replicability in their evolution. Adaptive paths were highly parallel for some strains, whereas others evolved distinct mutations across iterations of the experiment. Our data suggest that differences in replicability arise from mutational biases and variable fitness impacts of mutations across genetic backgrounds. Comparison of our results with genomic data from *M. tb* populations within hosts with tuberculosis (TB) revealed that several mutations associated with biofilm selection are also among the most common to emerge during natural infection. These biofilm-associated variants are not maintained in natural *M. tb* populations, suggesting that biofilm selection in our model mimics selection pressures that are transiently encountered by *M. tb* during specific phases of infection. Overall, these results support development of biofilm directed therapies for TB and demonstrate the importance of subtle genetic variation in shaping *M. tb* responses to changing selection pressures.

**Significance:** *Mycobacterium tuberculosis (M. tb),* the causative agent of tuberculosis (TB), a difficult-to-treat, persistent infection with high mortality. One cause of this persistence is the ability of *M. tb* to form biofilms, aggregated structures capable of resisting antibiotics and host defenses. Here, we used an evolutionary model to investigate mechanisms of *M. tb* biofilm formation. We found adaptation to biofilm growth to be affected by mutational biases and interactions among mutations. Our simple in vitro model appears to mimic aspects of natural infection, as identical mutations emerge in the laboratory under biofilm selection and within hosts with TB. These results expand our knowledge of *M. tb* biofilm development and inform our understanding of how this bacterium responds to novel selection pressures.

## Introduction

*Mycobacterium tuberculosis (M. tb)* is the causative agent of tuberculosis (TB), a disease that afflicted close to 11 million people and caused over a million deaths in 2021 (Bagcchi 2023). Active TB represents a small fraction of total infections, as *M. tb* can persist in a dormant state in the host for years and an estimated 2 billion people harbor latent *M. tb* infections (Flynn and Chan 2001; Bagcchi 2023). Latent M *tb* infections can transform into active TB, with immunocompromised patients (such as people living with HIV) at a significantly higher risk of developing active disease (Bagcchi 2023; Alsayed and Gunosewoyo 2023).

*M. tb* is capable of surviving in the host for long periods, despite immune defenses and antibiotic treatment (Wallis et al. 1999; Ojha et al. 2008; C. H. Liu, Liu, and Ge 2017; Richards et al. 2019). This tolerance has been linked with persister cells, a subpopulation of *M. tb* cells that are tolerant to high levels of antibiotics and other stressors (Joshi, Kandari, and Bhatnagar 2021). The ability of bacterial subpopulations to withstand antibiotic treatment necessitates prolonged combination therapy for active TB, which makes the disease difficult to control and to manage and in turn drives emergence of drug-resistant *M. tb* strains (Chiang, Centis, and Migliori 2010).

Growth as a biofilm has been identified as a mechanism by which *M. tb* can withstand antibiotics and other stressors (Ojha et al. 2008). Biofilms are increasingly understood to be an important form of bacterial growth across diverse species, where cells grow as a community encapsulated by an extracellular matrix (Costerton, Stewart, and Greenberg 1999). Biofilms confer tolerance to host defenses and antibiotics (Ciofu et al. 2022). Although TB is not traditionally thought of as a biofilm infection, *M. tb* is known to form aggregates in vitro, and those aggregates produce an extracellular matrix (Dubos and Davis 1946; Bacon et al. 2014). Additionally, *M. tb* aggregates have long been identified in human lung tissue (Canetti 1956; Walenty Nyka 1963; Nyka W 1967; W. Nyka and O’Neill 1970; W. Nyka 1977; Chakraborty et al. 2021).

Several in vitro models of *M. tb* biofilms have been developed, each of which demonstrates increased tolerance to antibiotics (Ojha et al. 2008; Ackart et al. 2014; Trivedi et al. 2016). The pellicle biofilm model developed by Ojha et al consists of *M. tb* grown on an air-liquid interface in minimal nutrient media, based on the finding of an acellular rim near necrotic tissue containing drug-tolerant *M. tb* microcolonies in guinea pig lung tissue (Lenaerts et al. 2007). Understanding *M. tb* biofilm formation can enable development of novel treatment strategies (F. Wang et al. 2013; Ackart et al. 2014; Richards et al. 2019; Chakraborty et al. 2021). Prior research in this area has targeted candidate biofilm genes via knockout, knockdown, and overexpression mutants and examined their impact on biofilm formation (Ojha et al. 2008; Pang Jennifer M. et al. 2012; Sambandan Dhinakaran et al. 2013; Wolff et al. 2015; Rastogi et al. 2017; Yang et al. 2017; Richards et al. 2019; Hegde 2020; Bharti et al. 2021; Chakraborty et al. 2021).

An alternative, and complementary method of investigating microbial traits is through experimental evolution. This technique involves growing a bacterial population under a specific selective pressure that drives adaptation toward a particular niche (Kawecki et al. 2012). This approach has the benefit of maintaining complex cellular networks intact, enabling the identification of subtle genetic changes associated with the phenotype under selection. Additionally, experimental evolution is unbiased regarding the selection of loci. There are multiple experimental evolution models of bacterial biofilm formation, which have been successful in identifying genetic determinants of biofilm growth as well as characterizing the complex communities within biofilms (Steenackers et al. 2016). For example, the well-established static microcosm model of *Pseudomonas fluorescens* has provided a wealth of insights into basic ecological and evolutionary processes as well as molecular mechanisms underlying important bacterial phenotypes (McDonald et al. 2009).

We developed an experimental evolution technique to study *M. tb* biofilm formation that is based on the pellicle biofilm model of growth at the air liquid interface (Smith et al. 2022). Our model uses unmodified clinical *M. tb* strains, which are more informative about *M. tb* in its natural environment than lab-adapted strains. An initial study passaging six closely related *M. tb* strains revealed convergent adaptation of bacterial populations under biofilm selection, i.e. bacterial genetic loci subject to repeated mutation in genetically distinct strains of *M. tb.* Here, we sought to investigate the overall replicability of *M. tb* adaptation by repeating the experiment, with triplicate lines for each ancestral genetic background. We found that the genetic background of each evolving population exerted strong impacts on replicability of its adaptation, likely as a result of selection on standing variation, mutational biases, and epistasis. We identified novel candidate loci underlying *M. tb* biofilm development, and comparisons of our experimental results with natural population data indicate that bacterial loci under selection in our experiments are also under selection within hosts with TB.

## Results

### Experimental design: replaying the tape of evolution

In previous work we investigated *M. tb* adaptation to pellicle growth by passaging six closely bacterial populations under selection for pellicle biofilm growth (Smith et al. 2022). Here, we resuscitated ancestral populations from these lineages and repeated the experiment in triplicate (Figure 1). We evolved the twelve populations in parallel with each other, imposing selection in the same manner as in the original study. In the framework of Blount et al, this represents both a parallel replay and historical difference experiment (Blount, Lenski, and Losos 2018). Historical difference refers to the genetic variants separating clinical isolates, which accumulated in *M. tb’s* natural environment. In our prior study, we found evidence suggesting that genetic differentiation among clinical isolates of *M. tb* affects their adaptation to a new environment. We sought to further investigate this pattern here, and to examine the importance of contingency in *M. tb* adaptation by ‘replaying the tape’ of evolution in the parallel replay experiments.

**Figure 1:**
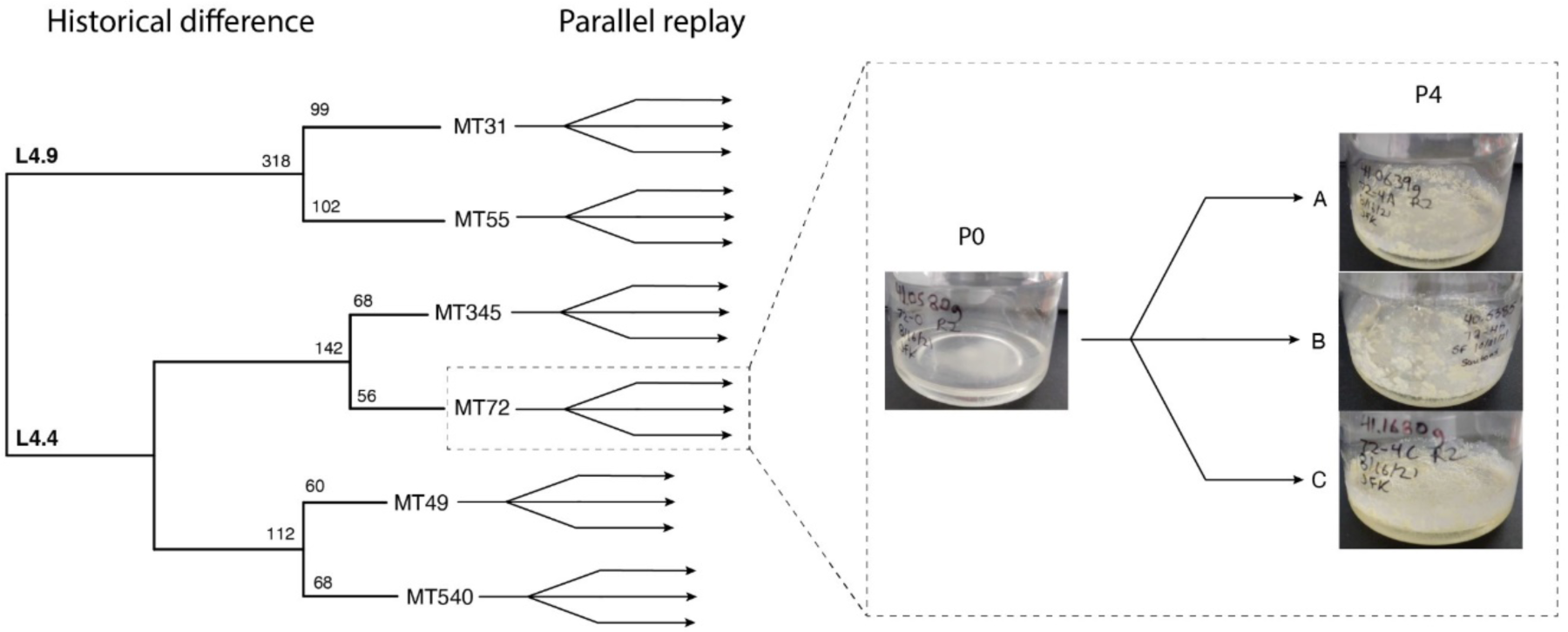
Experimental schematic for serial passaging of *M. tb* clinical isolates under selection for biofilm growth. Ancestral populations from six closely related clinical isolates (historical difference) were each split into three independently evolving lineages (parallel replay). The experimental design captures both historical difference (variation among clinical isolates that accumulated in a natural environment) and parallel replay (independently evolving replicates). Left: whole genome sequence phylogeny of ancestral populations used for original passaging experiment (Smith et al. 2022) shown with single-nucleotide polymorphism distances. Right: Example of gross morphology changes across passaging using pellicle photos from MT72 ancestor (Po) and all replicates after four passages (P4).

### Phenotypic changes in response to biofilm selection

We measured fitness in this system by weighing the pellicles, as in the original study (Smith et al. 2022). These data indicate that outcomes of the experiment were for the most part deterministic, in that fitness increased in the majority of populations as measured by wet weight (Figure 2). The exceptions are MT345 (all replicates) and MT55A, where we observed stable or decreasing wet weights following pellicle passaging.

**Figure 2:**
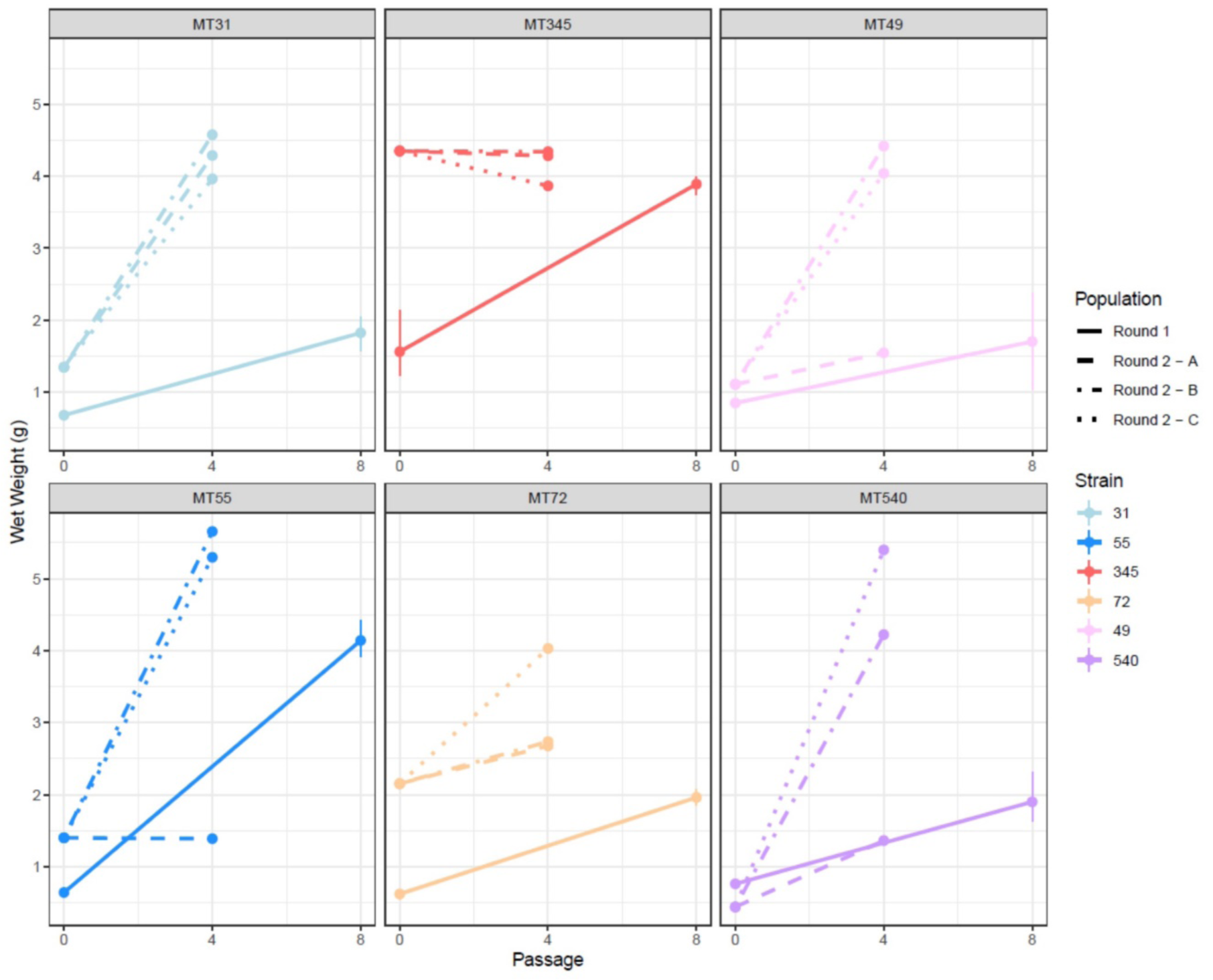
Wet weights of pellicle biofilms from twelve passaging lines of *M. tb.* Wet weights were measured as described in Smith et al 2022. Ancestral populations (passage 0) are shown from both rounds of the experiment, as well as after passaging under biofilm selection, for the original experiment (passage 8) and this round (passage 4). Ancestors from round 2 versus round 1 were subject to additional minor laboratory manipulation (additional freeze-thaw-grow cycle, see Methods for details). Pellicle wet weights for the most part increased following this non-specific laboratory manipulation, as well as after specific selection for biofilm growth. MT345A, B, and C as well as 55A did not increase wet weight following biofilm passaging.

Prior to the imposition of selection for biofilm growth, ancestral bacterial populations from the original study were frozen, and then thawed and re-grown for this study (see *Methods* for details). This minor manipulation in the laboratory appears to have affected biofilm fitness as ancestral populations from this round (hereafter, R2) in many cases exhibited higher wet weights than those from the original study (R1, Figure 2). Interestingly, we observed a possible effect of genetic background on susceptibility to minor laboratory manipulation. Pairs of populations from the same sub-lineage exhibited similar changes: R2 ancestral wet weights were substantially higher for L4.4.1.1, moderately higher for L4.9, and essentially unchanged for L4.4.1.2. For the most part, the fitness of evolved populations from this round of experiments was also higher than that of the first round (Figure 2).

We developed a qualitative classification system to describe the gross morphotypes of pellicle biofilms, which is shown in Figure S1 along with example photos. Pellicle morphotypes varied at baseline among the clinical strains, reflecting natural variation in this phenotype (Figure S2). We also observed differences between ancestral populations from the two rounds of passaging experiments, consistent with their ongoing adaptation during minor manipulation in the lab. This was especially marked for MT345, which also exhibited a large increase in wet weight (Figure 2). All populations evolved the same categorical phenotype in response to the imposition of a uniform selection pressure. This finding is a departure from the earlier iteration of the experiment, where we observed differences in pellicle phenotype among evolved populations (Smith et al. 2022). Similar to the first round, pellicles from evolved populations in these experiments were thicker than those of corresponding ancestral populations, with generally greater amounts of extracellular matrix (ECM) evident in SEM (Figure S2).

### Pre conditioning mutations emerge during laboratory manipulation

In addition to the phenotypic differences described above, sequencing data from R2 ancestral populations revealed genetic variants separating them from R1 ancestors (Table S1). In some cases the mutations appear to explain differences in phenotype between R1 and R2 ancestors. For example, the R2 ancestor of MT345 acquired a massive Hγokbp) duplication with an estimated frequency of 50% in the population (Figure 3). This duplication was maintained in all three replicates throughout pellicle passaging, suggesting it was advantageous under biofilm selection and potentially responsible for the large difference in wet weights of R1 vs R2 ancestors (Figure 2). The duplication persisted at a low frequency in one planktonically passaged population and was lost in the two others, consistent with biofilm-specific selection for this variant (Smith et al. 2022).

**Figure 3:**
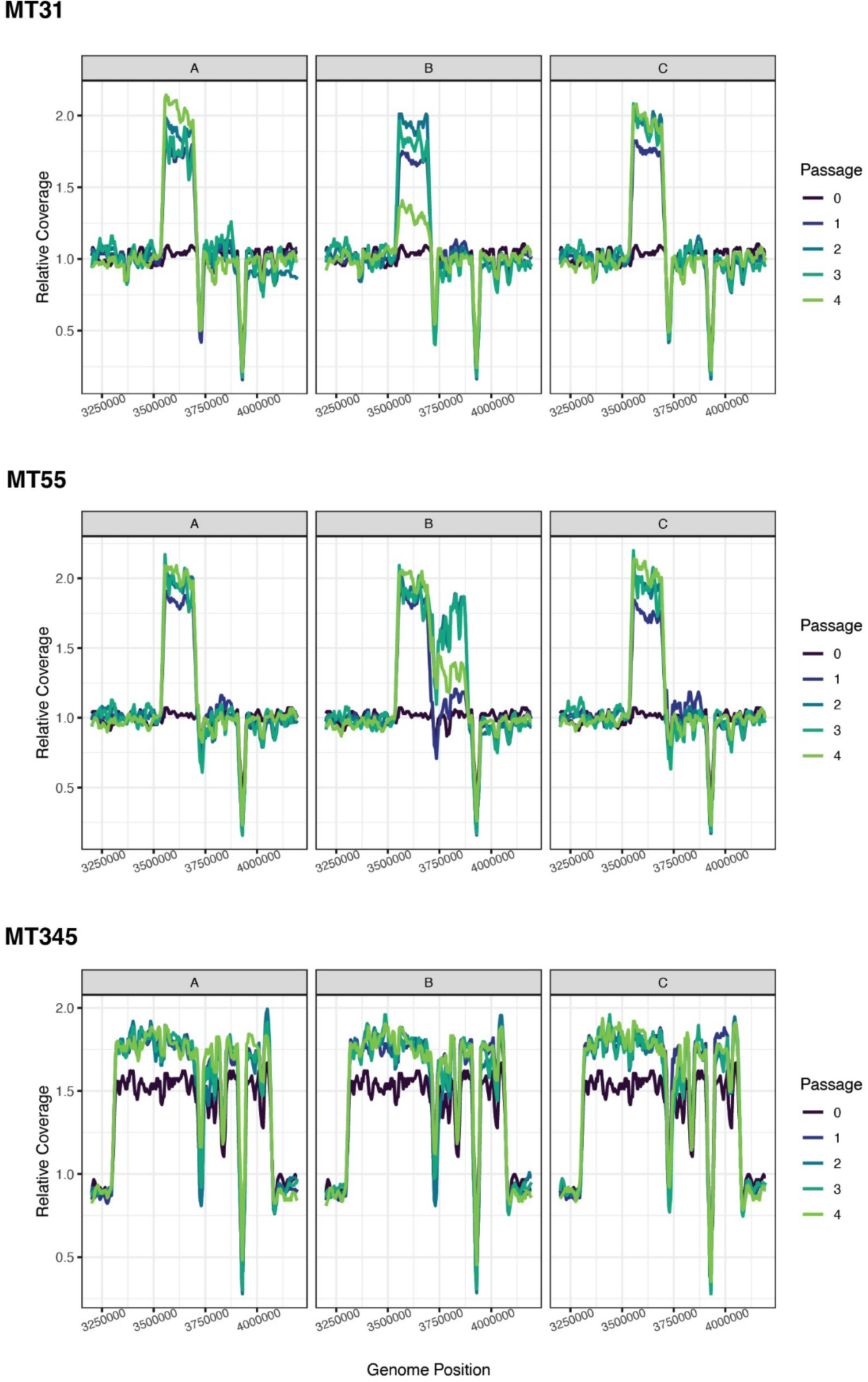
Large genomic duplications emerged repeatedly under biofilm selection. Plots show relative sequencing coverage calculated in sliding windows at positions between 3.25 - 4 Mbp. We identified similar genomic duplications in all six populations from the L4.9 sub-lineage (MT31 and MT55 strain backgrounds) and a massive duplication that was similarly maintained in all three replicate populations of L4.4.1.2 strain MT345. One L4.9 population (MT55B) initially evolved a larger duplication, which subsequently broke down to match the other L4. 9 populations.

A variant in *embR* segregating in the R1 ancestor of MT72 was lost in the R2 ancestor. The same variant was also lost during passaging in round 1 of the experiment. This suggests the variant could have a negative impact on fitness during pellicle growth and its loss may have contributed to the increased wet weight of the MT72 R2 ancestor. Two variants in MT31 were similarly lost during passaging in round 1, and in the R2 ancestor (Table S1).

Together, these observations show that *M. tb* evolves following minor routine manipulation in the lab, as shown by differences in pellicle phenotype, pellicle biomass, and genetic differentiation of ancestral populations. These data further suggest that this adaptation can affect forward evolution following imposition of a novel selection pressure.

### Identical mutations emerge under biofilm selection and within hosts with TB

Table 1 summarizes data for candidate loci subject to repeated mutation in association with selection for pellicle growth. Table S2 shows loci with allele frequency changes >=30% for each strain. Several novel candidate loci for pellicle adaptation emerged in these analyses including *mmaA1,* which has a role in mycolic acid modification, and *ilvC,* an essential gene that functions in valine and isoleucine biosynthesis. *lpdA* also plays a role in amino acid metabolism, in the interconversion of serine and glycine.

**Table 1:**
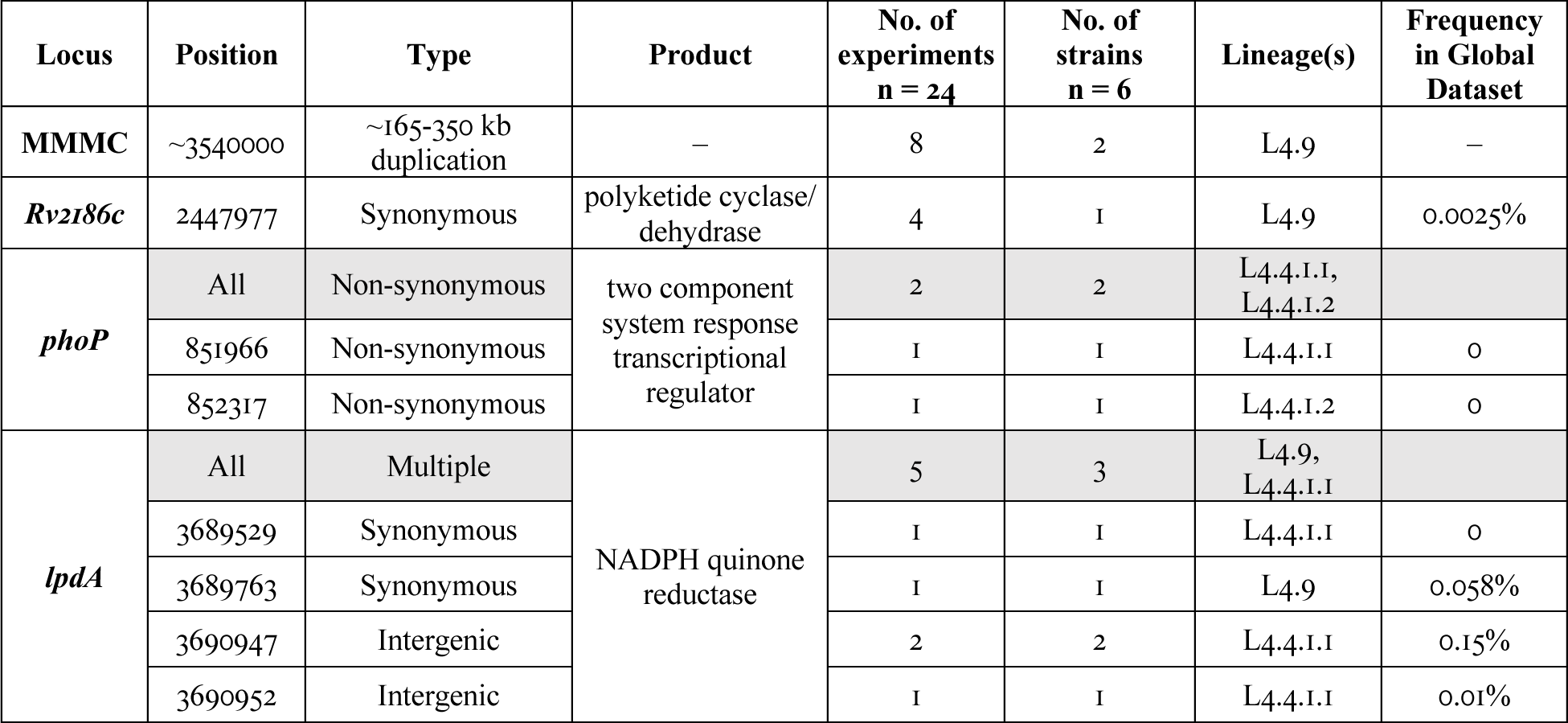
*M. tb* loci subject to recurrent mutation in response to biofilm selection. Table shows the number of passaging lines (out of 24), strain backgrounds (out of 6) and lineages (of 3) in which mutations were observed. For loci with multiple distinct mutations, summary data are provided and highlighted in grey. The frequency of each SNP in a natural population sample is shown (Global Dataset, see <u>Materials and Methods</u>).

We observed a striking degree of replicability across experiments, in some cases at the individual nucleotide level. The *lpdA* region was subject to recurrent mutation across the two rounds of the experiment, including an identical upstream mutation that occurred in both experiments. *lpdA* demonstrated a striking association with the MT540 background in particular: we observed mutations at *lpdA* in three of four passaging experiments with MT540. One of the R2 replicate populations, MT540-A, did not undergo mutation at *lpdA* and it achieved the lowest wet weight of any passaging experiment for this strain. In this population we identified a non-synonymous SNP in *phoP,* another locus subject to recurrent mutation.

We found previously that biofilm-associated mutations were rare or absent in a large sample of *M. tb* clinical isolates (Smith et al. 2022). Similar to our original study, the additional variants identified here were rare or absent from a natural population sample of close to 40,000 genomes (Table 1; Table S2, all variants). We discovered previously that one of the variants we identified upstream of *lpdA* in our original study swept to fixation within an individual host with TB (Trauner et al. 2017). Since our original study was published, Liu et al reported within-host variants from more than 50,000 samples (Q. Liu et al. 2022). Liu *et al* searched for evidence of positive selection at genic and intergenic regions by identifying loci subject to recurrent mutation within hosts, identified as unfixed mutations. The region upstream of *lpdA* had the largest number of unfixed mutation events of any intergenic segment in their analyses. Thus, it appears that the region subject to recurrent mutation under biofilm selection is also subject to recurrent mutation within hosts.

To further investigate similarities between biofilm selection and selection within hosts with TB, we examined patterns of variation at candidate loci from our experiments in the Liu dataset. We identified a striking pattern in these within-host data: the two synonymous *lpdA* variants that emerged under pellicle selection (Table 1) also emerged during TB infection; in fact, these two sites had by far the highest counts of unfixed mutations within the gene (Figure S3). The same was true of the intergenic region upstream of *lpdA’.* this region has the highest number of variants for any intergenic region in the within-host dataset, and the specific position that mutated repeatedly under biofilm selection had the highest mutation count in this intergenic region (Figure S3) and the third highest mutation count of any variant (genic or intergenic) in this large dataset. Mutations at these three sites appeared at a wide range of frequencies within hosts (Figure S3), with their overall distributions suggesting the mutations generally represent minority alleles. Integrating these observations, we hypothesize that *in vitro* selection for pellicle growth mimics one or more aspects of the environment within hosts with TB, and that this selection is at odds with evolutionary constraints encountered during transmission (antagonistic pleiotropy).

Two additional candidate biofilm loci were identified as potential sites of within-host selection: the region upstream of *ddlA* (Table S2) and *phoP* (Table 1). However, unlike the *lpdA* mutations described above, we did not identify any exact matches between biofilm-associated variants and unfixed mutations within hosts with TB.

### Parallelism and epistasis in emergence of genomic duplications under biofilm selection

Structural variants were prominent among mutations that emerged during pellicle passaging. For the two strains on the L4.9 background, there appears to be a deterministic relationship between pellicle passaging and emergence of a large duplication we term MMMC. This duplication emerged in every instance of passaging on this background, a total of eight times across the two rounds of experiments (Table 1; Figure 3). It did not emerge in planktonically passaged populations (Smith et al. 2022). A synonymous variant in *Rv2186c* also showed a consistent association with the MT31 background and with the duplication. This variant was segregating in the ancestral populations at 30 (R2)-39% (R1) frequency. It rose in frequency in all pellicle passaged populations, in association with emergence of MMMC, whereas it stayed stable in the planktonically passaged populations.

One replicate of MT31 followed the typical trajectory of MMMC emergence in association with a rise in frequency of the *Rv2186c* variant, but then subsequently developed a mutation at the *lpdA* locus. As the *lpdA* mutation rose in frequency, both MMMC and the *Rv2186c* variant declined.

### Multiple mechanisms of adaptation

Allelic trajectories, plotting variants that changed in frequency >= 30% over the course of the experiment are shown in Figure 4. These data implicate a variety of adaptive paths to pellicle growth. We observed likely clonal replacements/ hard selective sweeps *(lpdA,* MMMC, *mmaA1, ilvC),* selection on standing variation/ soft sweeps *(Rv2186c,* MT345 duplication), balancing selection *(Rv2186c,* possibly *phoP* vs incomplete sweep) and clonal interference *(lpdA* vs MMMC). For some populations, including strains with dramatic increases in wet weight (MT49-B, C) we did not identify any potentially explanatory variants.

**Figure 4:**
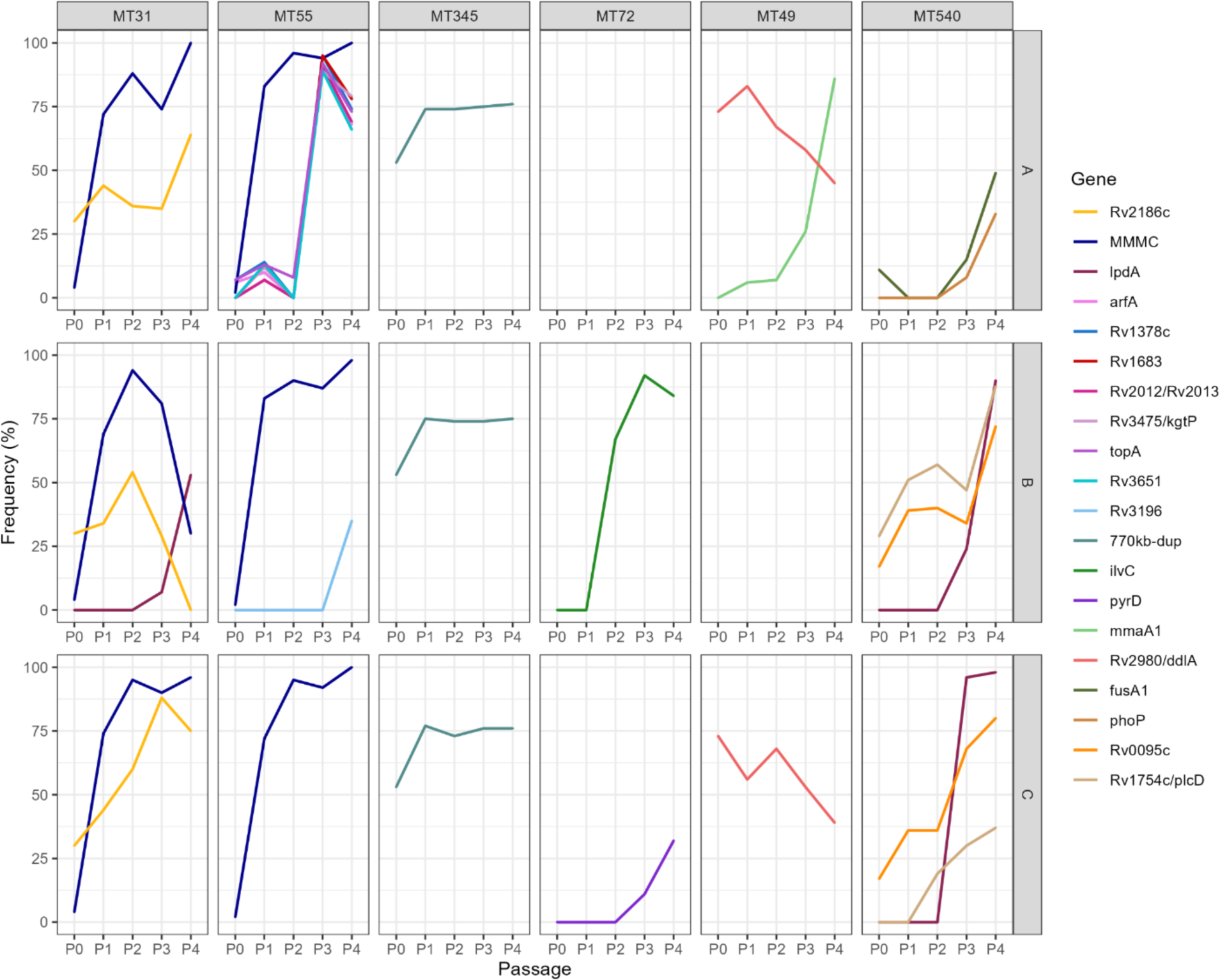
Multiple adaptive paths to biofilm growth. Allelic trajectories of mutations with >=30% changes in frequency during pellicle passaging, broken down by strain and replicate. Mutations that were also identified in planktonically passaged cultures are not shown. Mutation frequency data were calculated with Popoolation2 and plots were made using ggplot2 (see Materials and Methods).

### Tandem genomic duplications: frequent, reversible mutations

As with our earlier experiment (Smith et al. 2022), we generally identified a small number of mutations in association with pellicle adaptation. We did observe one exception, strain MT55-A. We identified a minority haplotype encoding a SNP in *topA* that was segregating at low frequencies in the ancestral population. It persisted during passaging and acquired four novel mutations by the first passage (Figure 4). This haplotype was below the limit of detection at the second passage but swept to fixation at the third passage. The MMMC duplication was already fixed at the second passage. By the fourth passage, the *topA* haplotype was again disentangled from the MMMC variant, as the MMMC remained fixed while the *topA* haplotype declined in frequency. Together, these observations suggest: 1) the MMMC variant segregated in the population both with and without the *topA* haplotype; 2) the MMMC variant is unstable, reverting at non negligible frequency and/or that small sub-populations lacking the duplication persist throughout passaging. Coverage plots of the MMMC region indicate that it was at 2X coverage from passage 2 onward (Figure 3), which could indicate that a duplication was fixed (or close to fixed), or that there were sub-populations where the region was duplicated multiple times and others that lacked any duplication. These data also suggest that the combination of the *topA* haplotype and the duplication has lower fitness than does MMMC alone as this combination did not persist in the population and final wet weights were close to the ancestral population for this replicate.

The allelic trajectories of the *Rv2186c* variant and MMMC on the MT31 background also imply that the duplication arose multiple times, both in association with the variant and without it, or that some bacteria encoded multiple copies of the duplicated region (Figure 4). Together, the observations of MT31 and MT55 point to a high degree of plasticity at the MMMC locus.

### Dynamics of co-circulating advantageous mutations

The allelic trajectories for MT31-B suggest that the MMMC and *lpdA* mutations were in competition, and that the *lpdA* mutation confers higher fitness, as it seemed to be overtaking the population (Figure 4). We performed two sets of experiments to test this hypothesis. In the first, we extended passaging of MT31-4B, carrying the population through an additional five passages under selection for pellicle growth. In the second set, we mixed MT31-4B and MT31-4C populations in a 50:50 ratio and passaged the mixed populations under selection for pellicle growth. In all cases, the allelic trajectories for *lpdA* and MMMC remained opposed, indicating they were not on the same background (Figure 5). Given the evidence for plasticity at the MMMC locus, this suggests the two mutations are incompatible. The *Rv2186c* variant was already lost at the outset of the extended passaging experiment. In the competition experiments, this variant followed the same trajectory as MMMC, as it has done in all other experiments on the MT31 background. Here we have evidence of different kinds of epistasis: additive or positive epistasis (MMMC & *Rv2186c* synonymous SNP (sSNP)) and sign epistasis (MMMC & *lpdA* sSNP). All three variants appear to be advantageous, based on their parallel evolution (MMMC *& lpdA)* or persistence across multiple experiments *(Rv2186c).* The combination of MMMC and *Rv2186c* sSNP appears to be more advantageous than either individual variant, which could result from additive effects on fitness or synergy (positive epistasis). By contrast, the combination of MMMC and *lpdA* sSNP appears to be deleterious (sign epistasis).

**Figure 5:**
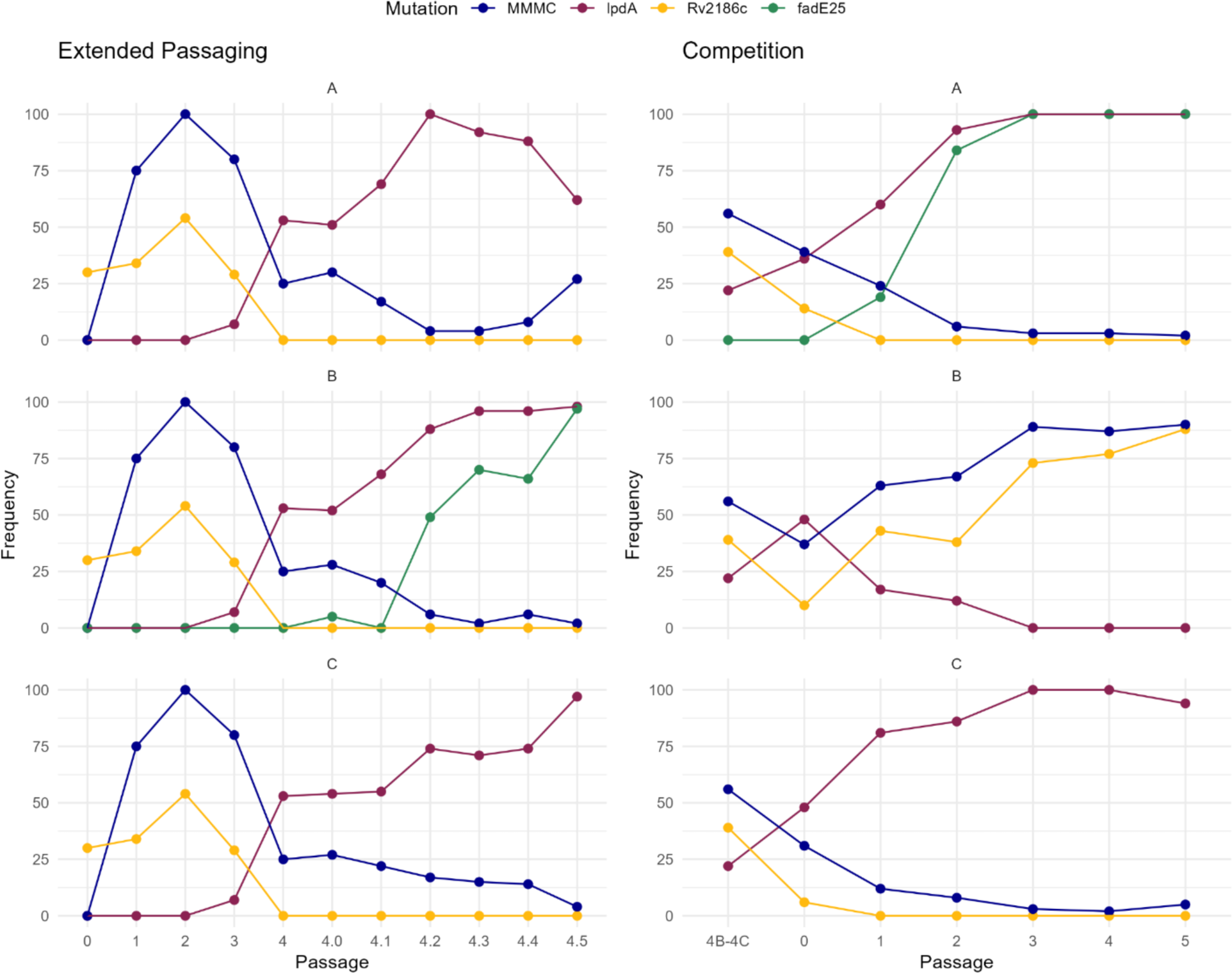
Dynamics of competing and cooperating mutations. Allele frequency trajectories are shown for synonymous SNPs in *lpdA* and *Rv2186c,* a non-synonymous SNP in *fadE25*, and the MMMC duplication. The *lpdA* SNP and MMMC were co-segregating at intermediate frequencies in passage 4 of MT31-B. Data from further passaging of this population (performed in triplicate, passages 4.0-4.5) are shown on the left. Passage 4.0 refers to the first biofilm grown from frozen stock. On the right, we show data from competition experiments in which equal proportions of population MT31-4B and MT31-4C were mixed (4B-4C) and passaged (performed in triplicate, passages 0-5). Passage 0 refers to the first biofilm grown from the mixed cultures. Allele frequency data for these experiments are shown in Table S3.

The *lpdA* variant landed at a higher frequency than MMMC in five of the six experiments (Figure 5), suggesting it confers a greater fitness benefit. However, the outcome of this competition was not deterministic, as shown by the one population in which the MMMC haplotype fixed to the exclusion of *lpdA* (MT31-4B4C-5B). Moreover, in one of the extended passaging experiments (MT31-4B-A), the frequency of the *lpdA* sSNP declined after fixing as the MMMC increased in frequency. A similar, more subtle pattern was seen in a competition lineage (MT31-4B4C-C). These data suggest that differences in fitness benefits of *lpdA* and MMMC mutations are moderate, and/or that the outcome of competition depends on additional contingencies.

We identified an additional site of parallel evolution in these experiments: identical non-synonymous mutations occurred in *fadE25* (E31Q), a probable acyl-CoA dehydrogenase involved in lipid degradation. The *fadE25* SNP was absent from both the global dataset of between-host variants, and the within-host dataset of Liu *et al*. This mutation swept to fixation along with the *lpdA* mutation (Figure 5), suggesting this combination is beneficial, similar to MMMC and the *Rv2186c* variant.

### Phenotypic impacts of gene dosage vary among genetic backgrounds

In our prior study (Smith et al. 2022), two strains from the L4.4.1.2 background underwent mutation at a transcription factor binding site (TFBS) upstream of *lpdA.* We found *lpdA* to be upregulated in strains with the TFBS mutations relative to their ancestral versions lacking these mutations. Transformation of H37RV with an additional copy of *lpdA* resulted in a significant increase in pellicle wet weight. Thus we hypothesized that the TFBS mutations resulted in de-repression of *lpdA* expression leading to an increase in pellicle growth. Here, we identified additional mutations at this locus, including synonymous mutations within the gene. In this round of the experiment, mutations occurred on the L4.9 background in addition to the L4.4.1.2 background. Our analyses of within-host data from TB patients demonstrated that mutations at this locus occurred on a variety of different genetic backgrounds (global lineages).

To further investigate interactions between genetic background, expression of *lpdA* and pellicle fitness, we transformed all ancestral (R2) strains from this study with *lpdA.* We also transformed the strains with a second copy of *glpD2,* which is adjacent to *lpdA* and resides within the same operon. Our transcriptomic data suggest these two genes are co-transcribed (Youngblom et al. 2024). Wet weights for the transformants are shown in Figure 6. We found the impact of transformation to vary dramatically among strains, with both positive, negative and neutral impacts on pellicle fitness. Strain MT540 could not be transformed with a second copy *of glpD2,* suggesting the mutation is lethal. L4.9 showed a potential impact of sub-lineage in that the impacts of transformation were concordant for the L4.9 strain pair: neither transformation with *lpdA* nor *glpD2* appeared to affect pellicle fitness (Figure 6). Impacts were not concordant for the other strain pairs in our study, nor were they concordant for transformation with the two genes. Together, these data suggest: 1) dosage of *lpdA* and *glpD2* can affect pellicle growth; 2) mutations at the *lpdA* locus affect pellicle growth but the mechanism is unlikely to be due solely to an increase in expression of this gene; 3) changes in the dosage of *lpdA* and *glpD2* have different impacts across genetic backgrounds.

**Figure 6:**
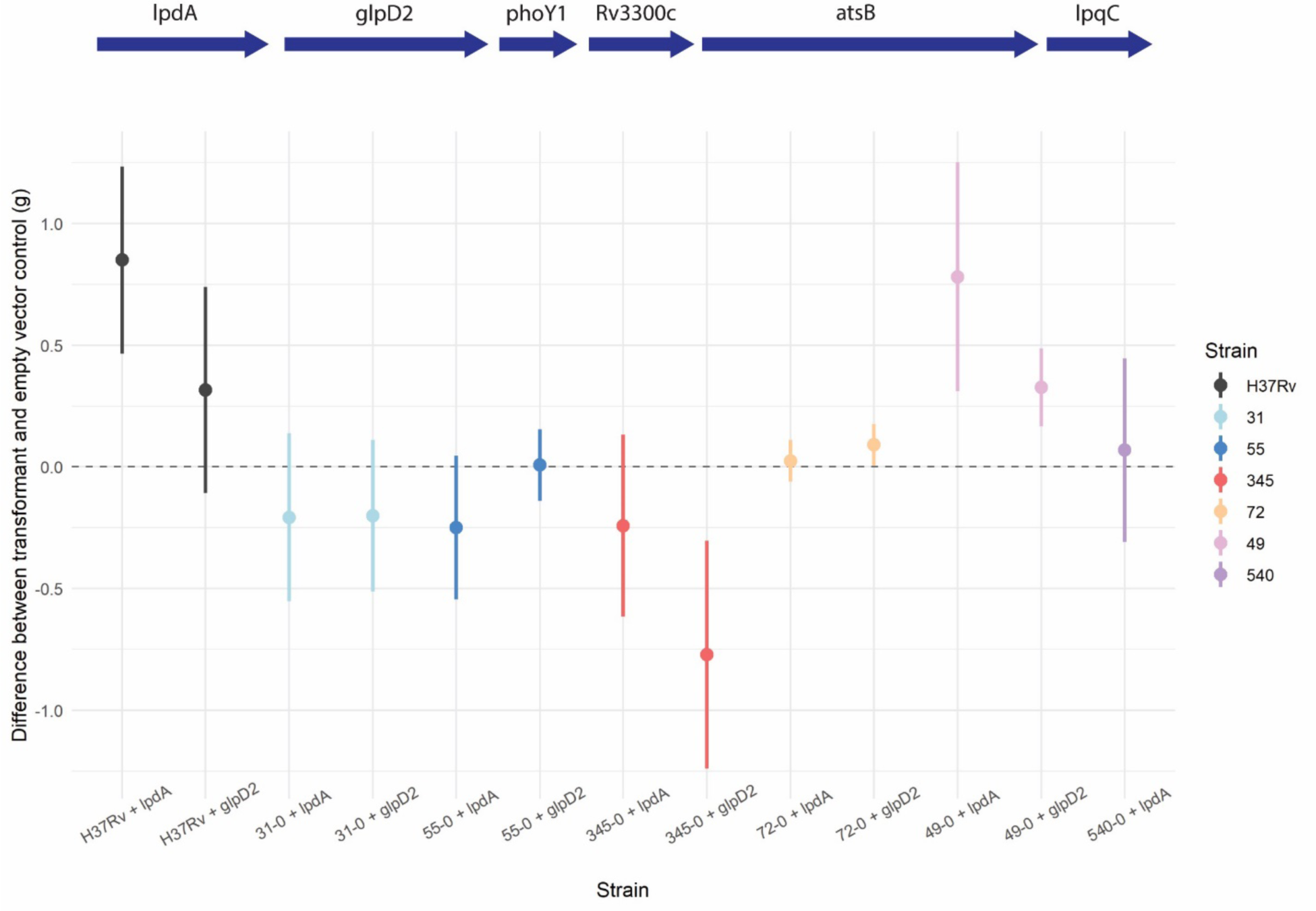
Phenotypic impacts of increased gene dosage vary among strains. Difference in pellicle wet weights between *lpdA* or *glpD2* transformants and the empty vector control are plotted for each strain. R2 ancestors were used for mutant construction, data from three replicates are shown. Differences between transformants and empty vector controls were highly significant (Kruskal-Wallis, p = 5.451e-05).

## Discussion

### Importance of large genomic duplications in M. tb adaptation

In our original experiment, we identified an ∼176kbp tandem duplication that emerged repeatedly and specifically under selection for pellicle growth (Smith et al. 2022). The duplication, which we term MMMC (Youngblom et al. 2024), emerged in association with the L4.9 sub-lineage. The current study shows that the association between L4.9 strains, pellicle selection and MMMC is highly replicable: the duplication emerged independently in every experiment with strains from this sub-lineage (total n=8, Table 1).

Tandem duplications are thought to be the most prevalent type of mutation across species (Reams et al. 2010) and prior research on model organisms has demonstrated relatively high rates of structural mutation during bacterial evolution in vitro (Schenk et al. 2022). We found previously that MMMC was maintained under biofilm selection over a prolonged period (up to 20 passages, ∼2 years), but not under planktonic conditions (Smith et al. 2022). Our results here indicate that the duplication occurred frequently, with repeated gain and loss events in our experimental *M. tb* populations (Figure 4). Together, these data suggest that MMMC was continually selected in a dynamic process as the variant was gained and lost. We hypothesize that the specific association of MMMC and L4.9 strains in our study is related to the location of flanking insertion elements, which typically mediate structural mutations in *M. tb* (Domenech et al. 2010; Shitikov et al. 2014; Smith et al. 2022). Mutational biases have been shown previously to produce parallelism in experimental evolution, and in some cases to interfere with fixation of the highest fitness mutations (Sackman et al. 2017; Horton et al. 2021). IS-mediated mutations are frequently observed in *M. tb* (Thabet and Souissi 2016) and we identified additional likely candidates in this experiment including a massive duplication in MT345 (Figure 3) and a transient MMMC duplication in one sample of MT49 (Youngblom et al. 2024).

We further hypothesize that the consistent association between L4.9 strains, biofilm selection, and the MMMC duplication reflects both the IS-driven mutational bias described above and fitness effects of the duplication that are specific to this genetic background. Smaller duplications (∼3kbp) have been shown to be key to development of a novel function (citrate utilization) during experimental evolution of *E. coli,* where the underlying mechanisms appear to be promoter capture and increased gene dosage (Blount et al. 2012). Dynamic gene amplification and loss have also been observed in natural bacterial populations under antibiotic pressure (Nicoloff et al. 2019). We do not know the mechanism by which MMMC affects biofilm growth: in a separate study we found the duplication does not simply double the dosage of genes within its borders, but rather has complex and in some cases dramatic impacts on expression of genes within and outside the duplicated region (Youngblom et al. 2024). Here, we identified a population (MT55B) with an initial duplication that extended beyond the borders of the MMMC: over the course of passaging, the duplication broke down such that it matched the MMMC (Figure 3). This suggests that selection for the duplication is precise, despite its impacts on multiple loci. We found previously that populations in our experiment exhibit different transcriptomic responses to biofilm growth, and that they converged on lineage-specific transcriptomic set points in response to pellicle selection (Youngblom et al. 2024). Here, we found perturbations in the dosage of a single gene to have distinct phenotypic impacts from strain to strain (Figure 6). Prior research has identified different effects of gene deletion and downregulation among *M. tb* from distinct lineages (Carey et al. 2018; Bosch et al. 2021). Our results demonstrate that the impacts of regulatory mutations can differ even among closely related strains, likely as a consequence of differences in underlying regulatory architecture. We hypothesize that precisely because the tandem duplications observed here affect expression of so many genes, they are advantageous under relatively narrow circumstances comprising specific combinations of environmental conditions and strain background. Further research into the *cis­*and ira/w-acting regulatory elements acting on, and found within, the duplication may provide further clues to its mechanism of action.

### A structural versus single nucleotide variant: competition, contingency, and incompatibility

Deletions have been predominant among the structural variants observed in prior studies of bacterial experimental evolution, in some cases resulting in a trend to genome size reduction over the course of the experiments (Raeside et al. 2014; Schenk et al. 2022). Large genomic deletions are well-documented in natural populations of *M. tb,* and in fact are prominent among mutations separating *Mycobacterium tuberculosis* Complex (MTBC) members with distinct host ranges (Pepperell 2022). We did not identify any large genomic deletions in our experiment, whereas genomic duplications were prominent among the mutations associated with pellicle adaptation. In experimental evolution of *E. coli,* where genomic deletions predominate, SNPs generally confer higher fitness benefits than structural variants (Raeside et al. 2014; Schenk et al. 2022).

In one of the passaging lines in which MMMC emerged and swept to high frequencies, we also identified *de novo* mutation at the *lpdA* locus (MT31-B; Figure 4). In our earlier experiment we found this locus to be positively selected, a finding that we replicated and extended in the present study (Table 1; (Smith et al. 2022)). The MT31 mutation is the fifth independent emergence of a single nucleotide variant at the *lpdA* locus under selection for pellicle growth. Allelic trajectories from MT31-B suggest that two advantageous mutations were competing. The *lpdA* mutation emerged later than the MMMC, consistent with higher rates of structural versus single nucleotide mutations as has been previously observed in experimental evolution of *E. coli* (Schenk et al. 2022). In Schenk et al’s study, estimated selection coefficients for structural mutations were also lower than those of SNPs. Our initial data suggested that the *lpdA* mutants could have relatively higher fitness, as the frequency of this allele rose while the frequency of MMMC decreased. To further test this hypothesis, we performed extended passaging of this line as well as competition experiments (see *Methods).* These experiments show that the outcome of competition between the *lpdA* allele and MMMC is not deterministic.

In five out of six passaging lines, the *lpdA* mutant initially outcompeted MMMC (Figure 5). However, in two of these experiments (extended passaging MT31-4B-A & competition MT31-4B4C-C), after an initial decline to undetectable the MMMC variant rose in frequency. Given that the duplication appears to be continually lost and re-formed, re-emergence of the MMMC in these populations likely represents repeated mutation.

The dynamics of the two competing mutations exhibit contingency, which could be due to their having similar impacts on fitness and/or a prominent role for genetic drift in the system. Biofilm populations typically exhibit substantial spatial heterogeneity, even in simplified in vitro models (Steenackers et al. 2016). We did not attempt to disrupt any existing spatial structure during passaging of our biofilms, as this would damage the biofilm structure, alter the imposed selection pressure, and it is technically infeasible to reliably produce an even suspension of cells from these pellicles. Thus, it is possible that drift arising from spatial structure affects the allele dynamics observed. Rising frequencies of MMMC toward the end of the experiments could, for example, reflect passaging of a biofilm segment in which the relative frequencies of MMMC and the *lpdA* mutation favored the duplication despite overall predominance of the *lpdA* mutation. An additional/alternative explanation for the observed dynamics is that the selective advantage of *lpdA* and MMMC mutations are similar. Whatever the relative role of genetic drift in this system, it did not impede recurrent sweeps of *de novo* mutations in this and earlier iterations of the experiment (Figure 4; Table 1; (Smith et al. 2022)).

Allelic trajectories of *lpdA* and MMMC mutations remained opposed in all cases, suggesting that the two mutations are incompatible. This could reflect sign epistasis, where the variants are individually advantageous but their combination is deleterious. A similar pattern has been observed previously during experimental evolution of *Saccharomyces cerevisiae* under both nutritional and antimicrobial pressure: first pass, large effect mutations that were individually advantageous were found to be deleterious in combination (Kvitek and Sherlock 2011; Ono, Gerstein, and Otto 2017). Incompatibility has been observed between mutations with both similar and different mechanisms of action (Kvitek and Sherlock 2011; Tenaillon et al. 2012).

Mutations may have additive effects on phenotype while exhibiting antagonistic interactions (sign epistasis) with respect to fitness, for example as a result of overshooting fitness-optimized levels of gene expression (Chou et al. 2014). Our transcriptomics study identified broad impacts of *lpdA* and MMMC mutations on the pellicle transcriptome (Youngblom et al. 2024) and the combination of these two mutations could similarly overshoot optimal dosage of key genes. Alternatively, the mutations could act by distinct, mutually incompatible mechanisms. Below we discuss another potential contributor to incompatibility of MMMC and *lpdA* mutations, which is global loss of evolvability following large duplication events.

### Impacts of genomic duplications on subsequent adaptation

Evolvability has been defined as the capacity to produce beneficial phenotypic change that is heritable (Payne and Wagner 2019). Prior research has shown that structural variants can reduce evolvability, for example by deleting genetic material capable of adaptive evolution (Consuegra et al. 2021; Schenk et al. 2022). Gene loss has however also been observed to increase evolvability (Helsen et al. 2020). A priori we might expect the duplications we observed to be less detrimental than deletions, or even to enhance evolvability. Gene duplication underlies the emergence of novel function in important gene families such as type VII secretion systems in Mycobacteria (Mortimer, Weber, and Pepperell 2017) and chromosomal duplications have been shown to increase the rate of adaptation in experimental evolution of yeast (Selmecki et al. 2015). However, in our experiment, we found evidence to suggest that large genomic duplications constrain *M. tb* evolvability.

MT345 developed a massive ∼770kbp duplication during routine laboratory manipulation, which was sustained during selection for biofilm growth (Figure 3, Table 1). The duplication was associated with a large increase in wet weight in the ancestral population for this round of experiments (Figure 2). However, there was no measurable adaptation in bacterial populations with the duplication under pellicle selection: we did not detect any additional mutations, and fitness remained stable or decreased (Figure 2; Table S2). This could represent a form of diminishing returns epistasis, where positive effects of subsequent mutations are reduced on an increasingly fit genetic background. This phenomenon is well described in the experimental evolution literature, where it has been demonstrated as reduced fitness increments over time, reduced adaptive capacity in pre-adapted populations, and minimal incremental fitness benefits of mutations engineered into genetic backgrounds containing advantageous mutations (Barrick et al. 2010; Chou et al. 2011; Khan et al. 2011; Wiser, Ribeck, and Lenski 2013; Kryazhimskiy et al. 2014; Y. Wang et al. 2016; Wünsche et al. 2017). A prior study also identified an important role for ancestral fitness in shaping evolvability under a novel selection pressure, with highly fit strains of *S. cerevisiae* exhibiting lower evolvability (Jerison et al. 2017). However, we did not identify a similar pattern here, as highly fit ancestral populations exhibited large fitness gains across our two studies (Figure 2). Rather we observed reduced evolvability in association with genomic duplications, which we hypothesize arise from their global regulatory impacts, rather than generic fitness maxima.

Similar to MT345 with the massive duplication, populations with the MMMC duplication appeared constrained with respect to further evolvability. We found previously that the duplication was stable over years of passaging and did not observe any further sweeps on this background (Smith et al. 2022). Populations with the duplication behaved similarly in this round of experiments. For MT31, we did not observe additional mutations on the MMMC background, even during extended passaging (Figure 5; Table S3). It’s also notable that we observed clonal interference between the MMMC mutation and an *lpdA* mutation in MT31-B, suggesting that a competing mutation is more likely to emerge than a modification of the duplication. The high rate of structural mutations, and their reversibility, may enable emergence of competing mutations with greater evolvability as observed here.

For MMMC on the MT55 background, we identified additional mutations on the MMMC background in one of the replicate populations (MT55-A), and these appeared to decrease fitness. One of these variants is a synonymous SNP in DNA topoisomerase I *(topA). topA* variants have been found to be advantageous in several populations of the LTEE and have been associated with a hypermutator phenotype characterized by an increased frequency of tandem repeats (Crozat et al. 2005; Bachar et al. 2020). Whatever the individual impact of these additional variants may have in our *M. tb* populations, they appear to have been detrimental to fitness in combination with the MMMC. MT55-A had the lowest wet weight of any replicate, unchanged from its ancestors (Figure 2).

Observations of MMMC contrast with the *lpdA* locus. In extended passaging and competition experiments, we identified parallel, identical non-synonymous SNPs in *fadE25.* This SNP twice swept to fixation along with the *lpdA* sSNP (Figure 5). *FadE22* encodes an acyl-coA dehydrogenase. In the first round of our experiment, we also identified a sweep at an acyl-coA dehydrogenase *gene,fadE22,* again in conjunction with a sweep at *lpdA* (Smith et al. 2022). These data collectively implicate *fadE22* and *fadE25,* and perhaps acyl-coA dehydrogenase genes in general, in pellicle growth. They are also consistent with positive interactions between *fadE22/fadE25* mutations and SNPs at the *lpdA* locus. Lastly, they demonstrate that unlike genomic duplications in our experiments, *lpdA* mutations do not hinder evolvability.

Synthesizing our observations, large tandem duplications appear to enable rapid shifts in transcriptomic space that are advantageous to *M. tb* under a narrow set of conditions. We hypothesize that the transcriptome following duplication is relatively fragile, and thus there are fewer fitness-enhancing mutations available to these populations than to those carrying SNPs with more precise impacts on phenotype. Differences in evolvability following single nucleotide-versus structural mutations have been observed previously for *Pseudomonas aeruginosa* (Gifford, Toll-Riera, and MacLean 2016). Further research may provide insight into more and less robust transcriptomic states for *M. tb*.

### Standing genetic variation shapes evolution in a novel environment

The bacterial populations in our study were genetically differentiated at baseline, comprising six clinical strains from two sub-lineages (Smith et al. 2022). We found here that the strains underwent further mutation and differentiation during minor manipulations in the laboratory, prior to the imposition of biofilm selection (Table S1). Both types of genetic variation affected evolution under a novel selection pressure. As discussed above, the massive duplication that emerged in the MT345 ancestor appeared to both increase biofilm fitness and inhibit further evolvability. For most of the other strains, biofilm fitness increased to a greater extent than it did in the first round of passaging, suggesting that mutations acquired between experiments increased evolvability (Figure 2). In addition to the larger delta in biofilm fitness observed here, with the exception of MT345, the maximum wet weight achieved in this round of experiments was greater than in the initial round. *M. tb* populations remained highly dynamic as they were manipulated in the laboratory: genomic data shows loss of standing genetic variation, and emergence of *de novo* SNPs and structural variants during a single round of freezing and re-growth (Table S1). Some of the changes observed following this minor, non-specific manipulation recapitulate events observed in the first round of pellicle selection experiments: specifically, loss of segregating variants in *embR, acg,* and *nth* (Table S1). Novel variants that emerged in the R2 ancestors were also maintained in our experimental populations, suggesting that they were beneficial. Based on these observations, we hypothesize that variants emerging during non-specific laboratory manipulation can act to potentiate evolvability under pellicle selection. Potentiating variants have been shown to play key roles in emergence of complex traits during experimental evolution of bacteria, including citrate utilization in *E. coli* (Blount et al. 2012; Quandt et al. 2014; 2015; Douglas et al. 2017).

The effects of older genetic variants - those defining the strains and sub-lineages - were evident in the degree of parallelism observed across experiments. As discussed above, strains belonging to the L4.9 lineage exhibited a high degree of parallelism, evolving an identical mutation (MMMC) across all eight iterations of the experiment. Further, in every instance of pellicle selection on the MT31 background in this and our earlier experiment, MMMC has emerged in association with a synonymous SNP in *Rv2186c* that was segregating in the ancestral population (Table 1). MMMC and the *Rv2186c* sSNP have in all cases followed parallel trajectories during pellicle passaging (Figure 4), rising together in this and our earlier study. Frequencies of the *Rv2186c* sSNP remained stable in planktonically passaged populations. Thus, positive interactions between the MT31 background, the *Rv2186c* sSNP, and MMMC appear to be responsible for a high degree of parallelism in MT31 adaptation under pellicle selection. Parallelism was also evident on the MT540 background, with three out of four independent passaging experiments leading to emergence of mutations at the *lpdA* locus (Table 1). Parallelism was less obvious for the other strains in our study. Unlike MT31, MT55 and MT540, neither MT72, MT345 nor MT49 underwent parallel, *de novo* mutations under pellicle selection [(Smith et al. 2022) Table S2].

These observations suggest that adaptive landscapes differ among strains of *M. tb*. The adaptive landscape is a conceptual model of relationships among genotypes and phenotypes, where positions on a grid denote mutational combinations and height represents fitness [reviewed in (De Visser and Krug 2014)]. Fitness landscapes affect the amount of parallelism observed during adaptation, with ‘smooth’ landscapes associated with replicable evolution (Blount, Lenski, and Losos 2018). Negative interactions among mutations, particularly sign epistasis, can create fitness valleys and ‘rugged’ landscapes, which impede evolvability (Poelwijk et al. 2011). Sign epistasis is also associated with contingent outcomes, where newly advantageous mutations are deleterious on some backgrounds, forcing sub-populations to adopt distinct mutational paths (Carroll, Lee, and Marx 2014). We have both direct and indirect evidence of sign epistasis in our experimental system: transformation data demonstrate that changes in gene dosage are advantageous on some backgrounds and deleterious on others (Figure 6) and dynamics of MMMC and *lpdA* suggest they are incompatible with each other (Figure 5). Tandem duplications appeared to be broadly incompatible with new mutations, suggesting that they could impede access to a fitness optimum. We hypothesize that adaptive landscapes of the strains in our study vary as a function of both negative and positive epistasis. We also see an important role for mutational supply in shaping adaptive trajectories, evidenced in the prominence of standing genetic variation and high-rate mutations (duplications) in our experiments. These phenomena are in turn shaped by genetic background, in the form of ancestral within-population variation and background-specific mutational biases.

Our observations suggest that subtle genetic differences found in natural and experimental populations of *M. tb* have pervasive impacts on adaptation to novel environments. Within its natural environment, populations of *M. tb* encounter distinct selection pressures over time, for example during airborne transmission, the initial colonization of alveolar spaces, tissue invasion, spread, growth within a cavity and re-emergence into the airways. As micro-environments within hosts are encountered repeatedly across space and time, *M. tb* ‘replays the tape’ of evolution (Blount, Lenski, and Losos 2018). Our results suggest that for some *M. tb* genotypes, this will result in emergence of an array of advantageous mutations whereas for others the same mutations will repeatedly emerge.

Detailed studies of within-host evolution have demonstrated that *M. tb* infections comprise multiple, spatially separated sub-populations evolving autonomously (Lieberman et al. 2016; Martin et al. 2017; Moreno-Molina et al. 2021). Contingencies in *M. tb* evolution can produce more complex, heterogenous populations when these previously isolated pulmonary sub-populations mix in the airways, even when bacteria emerge from similar niches. Studies of *M. tb* within sputum have demonstrated such diversity, as sputum harbors genetically and phenotypically diverse bacteria (Garton et al. 2008; Séraphin et al. 2019; Nimmo et al. 2019; Shockey, Dabney, and Pepperell 2019). These diverse sub-populations may contribute to the robustness of the metapopulation as *M. tb* shifts to the new environment within airways.

Environmental fluctuations across niches within hosts are additional potential contributors to the robustness of *M. tb* metapopulations as mutational combinations that are incompatible in one environment may be advantageous in another; fluctuating environments can enable populations to overcome sign epistasis (de Vos et al. 2015; Ono, Gerstein, and Otto 2017). Such fluctuations may also enable acquisition of conditioning mutations as we observed here following non-specific manipulation of bacteria in the laboratory (Table S1).

### M. tb locus under selection for pellicle growth and within hosts with TB

In our prior study, we identified convergent evolution at a transcription factor binding site (TFBS) upstream of *lpdA* in two strains belonging to the L4.4.1.1 sub-lineage, MT49 and MT540 (Smith et al. 2022). The intergenic SNPs are associated with upregulation of *lpdA,* and transformation of H37RV with an additional copy of *lpdA* results in increased pellicle wet weight (Smith et al. 2022; Youngblom et al. 2024). Here we found further support for an association between *lpdA* and pellicle growth, as we observed repeated mutations at this locus under pellicle selection: twice on the MT540 background, and once on the MT31 background (Table 1). Our transcriptomic data indicate that genes within the *lpdA* operon are not co-transcribed, and are differentially regulated in pellicle versus planktonic growth (Youngblom et al. 2024). The present study reveals further complexity, in that the fitness impacts of transformation with *lpdA* and another gene within the operon, *glpD2,* are different from each other and vary dramatically from strain to strain, with positive, negative and neutral effects of transformation (Figure 6). We hypothesize that the strong association between MT540 and *lpdA* mutations (Table 1) reflects distinct features of its adaptive landscape, namely that that the relative fitness benefits of *lpdA* mutations are higher on the MT540 background than on other strains in our sample. Unlike L4.9 and MMMC, there is no obvious role for mutational bias in parallel evolution of MT540, as independently evolved populations developed different mutations at the *lpdA* locus (Table 1). In support of our hypothesis, we found that MT540 and MT345 evolved similar mutations in *phoP* but the *phoP* mutant had dramatically higher wet weight on the MT345 background, whereas a similar mutation conferred modest benefits on the MT540 background albeit at a lower frequency (Figure 2).

In this replay of our earlier experiment, we observed synonymous genic variants within *lpdA,* in addition to an intergenic SNP at an identical site as in the initial experiment. Given their proximity, it seems likely that the synonymous and intergenic SNPs act by a similar mechanism to increase biofilm fitness. Synonymous variants were previously thought to be neutral, but there is now increasing recognition that they can be under selection and important for bacterial adaptation (Lebeuf-Taylor et al. 2019). The mechanistic basis of selection on synonymous sites is an active area of research; available data suggest that in many cases variants affect phenotypes via gene dosage, e.g. by creating novel promoter sites, or affecting transcriptional or translational dynamics (Bailey, Alonso Morales, and Kassen 2021). Synonymous variation has also been shown to shape mutational biases (Horton et al. 2021). Synthesizing our observations across studies, we hypothesize that the intergenic and synonymous variants affect biofilm fitness using a similar regulatory mechanism, which affects dosage of genes within the *lpdA* operon and outside of it.

We found previously that the two intergenic SNPs upstream of *lpdA* are exceedingly rare in a sample of close to 40,000 consensus sequences of *M. tb* clinical isolates (Smith et al. 2022). However, in reviewing published data, we also discovered that one of the variants swept to fixation within an individual with TB (Trauner et al. 2017). Based on these observations we hypothesized that variants at the *lpdA* locus are transiently selected during infection. More recently, a large survey of within-host adaptation was published, comprising analyses of *M. tb* genomic data from more than 50,000 samples (Q. Liu et al. 2022). The purpose of Liu et al’s analyses was to identify intergenic regions (IGRs) and genes under positive selection within hosts; the IGR upstream of *lpdA* is among the regions that they identified with evidence of positive selection. In our review of the data from this study we found that the specific site subject to recurrent mutation in our experiments also underwent repeated mutation within hosts with TB - in fact, it had the third highest mutation count of any site in the genome (Figure S3) - and that the same was true of the synonymous sites within *lpdA* (Figure S3). In consensus sequences of *M. tb,* which collapse the diversity within individual hosts, these variants are exceedingly rare or completely absent (Table 1). These additional observations provide strong support for our hypothesis: identical variants are selected both during natural TB infection and under pressure for pellicle growth. This parallels studies of other bacterial infections such as *Staphylococcus aureus, Pseudomonas aeruginosa,* and *Burkholderia* spp, where the same loci appear to be under selection within hosts and during biofilm passaging in the laboratory [reviewed in (Steenackers et al. 2016). For M *tb,* the absence of pellicle biofilm-associated variants from consensus sequences suggests that these variants are deleterious during one or more phases of the bacterium’s natural life cycle. We favor that these variants are detrimental to transmission and/ or the initial phases of infection. Given the pleiotropic impacts of biofilm-associated mutations (Smith et al. 2022), it is not clear at this point whether selection within hosts is specific to biofilm growth, e.g. enhanced production of extracellular matrix, or other phenotypes.

### Summary

In this parallel replay experiment, we sought to evaluate the role of contingency in *M. tb* adaptation to a new environment. We discovered pervasive impacts of genetic background on the predictability of evolution in our system. Mutational supply played an important role in shaping adaptive walks, and genetic background exerted an impact on mutational supply via standing genetic variation present at baseline, as well as via background-specific mutational biases. Our results point to the importance of both positive and negative epistasis in shaping *M. tb* adaptation, with similar mutations having different impacts on different genetic backgrounds. Baseline fitness did not appear to affect evolvability in our experiments, but we did find evidence that certain mutation types could constrain forward evolvability. The complexity of within host environments, which are spatially structured and fluctuating, may protect against such evolutionary dead ends; even in the simplified environment studied here, we observed emergence of competing advantageous mutations. Some of the mutations observed in our experiments are also transiently selected within hosts with TB, suggesting that this simple in vitro system mimics aspects of natural infection.

## Materials and Methods

Briefly, six *M. tb* clinical strains were evolved under selective pressure to grow as pellicle biofilms, as described previously (Smith et al. 2022). In this iteration of the experiment, we evolved each clinical strain in triplicate, to examine replicability of adaptation under biofilm selection. Scripts used in the analysis of these sequences are available at https://github.com/pepperell-lab. A detailed account of materials and methods used in this study, along with data reported in the study, are available in **SI Appendix 2**.

## Acknowledgments

This work was supported by NIH NIAID R01AI113287 to CSP, NSF DGE-1747503 to MAY, NIH NIAID T32AI055397 to JFK, NIH NIAID T32AI007635 to JFK.

The authors wish to thank Dr Sarah Fortune for providing the overexpression plasmid and for assistance with transformation protocols.

## Notes

https://github.com/pepperell-lab

### Competing Interest Statement

The authors have declared no competing interest.

