## Supplementary Data 1 for "Contingencies in biofilm adaptation of *Mycobacterium tuberculosis*"

SI Appendix 1

**Figure S1:**

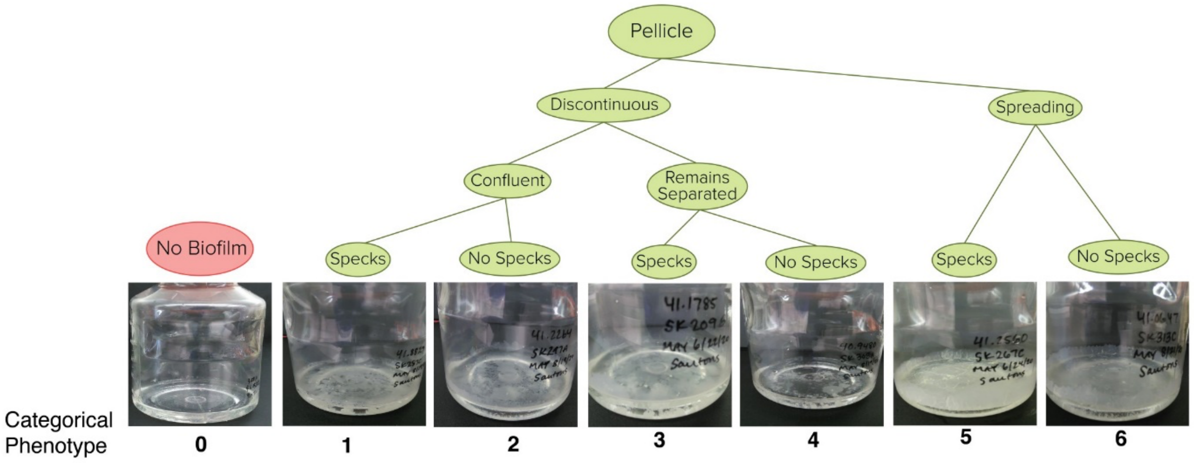

**Categorical phenotyping of *M. tb* pellicles.** We used a decision tree (green and red flowchart) to develop a categorical phenotyping scheme for *M. tb* pellicles. Each phenotype is denoted with a number from 0-6, which is shown below the example photos of each pellicle type.

**Figure S2:**

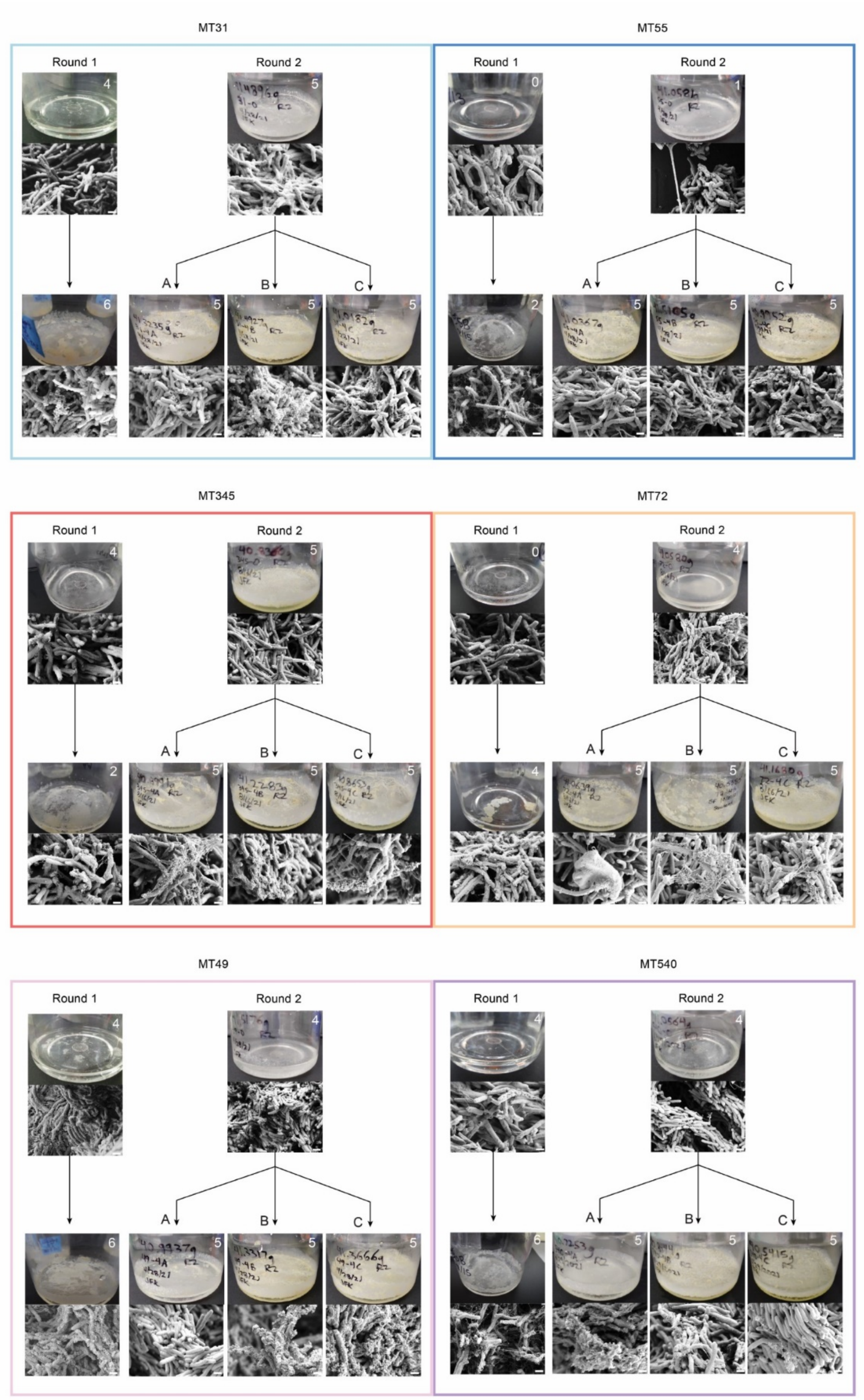
MT31
Round 1
Round 2
A
B
C
MT55
Round 1
Round 2
A
B
C
MT345
Round 1
Round 2
A
B
C
MT72
Round 1
Round 2
A
B
C
MT49
Round 1
Round 2
A
B
C
MT540
Round 1
Round 2
A
B
C

**Morphotypes of *M. tb* pellicle biofilms.** Photos showing gross morphotypes and SEMs for R1 and R2 ancestors, as well as R1 evolved populations (passage 8) and R2 evolved populations (passage 4). Images arranged by sub-lineage. Top: L4.9, middle: L4.4.1.2, bottom: L4.4.1.1. Numbers in upper right corner of pellicle photos refer to categorical phenotyping classification used to describe pellicle morphotypes (Figure S1). SEM micrographs are shown for the same passage points. Gross morphology and scanning electron micrographs were taken after 5-7 weeks of pellicle growth. Scale bars in lower right side of SEM images = 1 μm. Pellicle morphotypes varied among ancestral populations and changed following minor laboratory manipulation as evidenced by differences in R1 versus R2 ancestors. Pellicles of R2 ancestors were generally more robust in appearance, consistent with wet weight data; this is particularly marked for MT345. The R2 ancestor of MT72 exhibited more ECM than its R1 ancestor. ECM generally increased in response to passaging, across populations, and all populations in R2 converged on the same categorical morphotype (5) in response to pellicle passaging.

**Table S1:**

| Strain | Position | R1 Ancestor Freq. | R2 Ancestor Freq. | Type | Mutation | Gene | Product | Frequency in Global Dataset | Notes |
|---|---|---|---|---|---|---|---|---|---|
| **MT31** | 2279599 | 41 | 0 | Synonymous | D157D (GAT-GAC) | *acg* | NAD(P)H nitroreductase | 0 | identical variant fell out after R1 passaging |
| | 4359135 | 0 | 41 | Synonymous | P216P (CCA-CCG) | *espK* | ESX secreted effector | 0 | |
| | 4115505 | 32 | 0 | Synonymous | P116P (CCG-CCT) | *nth* | endonuclease III | 0.01% | identical variant fell out after R1 passaging |
| **MT55** | 2439519 | 43 | 0 | Synonymous | R143R (CGC-CGT) | *Rv2177c* | transposase | 0 | |
| **MT345** | 3310000 | 0 | ~50 | ~770 kb duplication | – | *lppX* – *Rv3639c* | – | | |
| **MT72** | 1416232 | 30 | 0 | Synonymous | C372C (TGT-TGC) | *embR* | transcriptional regulator | 0 | identical variant fell out after R1 passaging |
| | 1474778 | 0 | 71 | Non-coding | pos. 1121/3138 (C-G) | *rrl* | 23S ribosomal RNA | 0 | |

**Mutations identified following resuscitation of ancestral populations.** These mutations appeared in ancestral populations following minor laboratory manipulation at the initiation of the experiment, prior to the imposition of specific selection for biofilm growth.

### Table S2:

## MT31

| Replicate | Position | Frequency at passage 0 | Frequency at passage 4 | Type | Mutation | Gene | Product | Frequency in Global Dataset | Found in planktonically passaged populations |
|---|---|---|---|---|---|---|---|---|---|
| A | 2447977 | 30 | 64 | Synonymous | I6I (ATC-ATT) | *Rv2186c* | hypothetical protein | 0.0025% | |
| A | 3540000 | 0 | ~100 | 165kb duplication | - | *hpx – sdhA* | - | | |
| B | 2447977 | 30 | 0 | Synonymous | I6I (ATC-ATT) | *Rv2186c* | hypothetical protein | 0.0025% | |
| B | 3689763 | 0 | 53 | Synonymous | A392A (GCC-GCT) | *lpdA* | NAD(P)H quinone reductase | 0.058% | |
| B | 3540000 | 0 | ~25 | 165kb duplication | - | *hpx – sdhA* | - | | |
| C | 2447977 | 30 | 75 | Synonymous | I6I (ATC-ATT) | *Rv2186c* | hypothetical protein | 0.0025% | |
| C | 3540000 | 0 | ~100 | 165kb duplication | - | *hpx – sdhA* | - | | |

## MT55

| Replicate | Position | Frequency at passage 0 | Frequency at passage 4 | Type | Mutation | Gene | Product | Frequency in Global Dataset | Found in planktonically passaged populations |
|---|---|---|---|---|---|---|---|---|---|
| A | 1003702 | 6 | 68 | Synonymous | T297T (ACC-ACT) | *arfA* | peptidoglycan-binding protein | 0 | |
| A | 1551738 | 7 | 74 | Synonymous | P306P (CCG-CCA) | *Rv1378c* | hypothetical protein | 0.0025% | |
| A | 1909116 | 0 | 78 | Non-synonymous | R508Q (CGG-CAG) | *Rv1683* | long-chain acyl-CoA synthase & lipase | 0 | |
| A | 2260229 | 0 | 69 | Intergenic | +903/-436 (C-T) | *Rv2012 / Rv2013* | hypothetical protein / transposase | 0 | |
| A | 3540000 | 0 | ~100 | 165kb duplication | - | *hpx – sdhA* | - | | |
| A | 3892191 | 0 | 79 | Intergenic | +100/+180 (C-T) | *Rv3475 / kgtP* | transposase / dicarboxylate transport protein | 0 | |
| A | 4085524 | 7 | 73 | Synonymous | V578V (GTC-GTT) | *topA* | DNA topoisomerase I | 0.0025% | |
| A | 4091984 | 0 | 66 | Synonymous | A48A (GCC-GCT) | *Rv3651* | hypothetical protein | 0 | |
| B | 3540000 | 0 | ~100 | 165-350kb duplication | - | *hpx – bpoA* | - | | |
| B | 3566678 | 0 | 35 | Synonymous | C297C (TGT-->TGC) | *Rv3196* | hypothetical protein | 0 | |
| C | 3540000 | 0 | ~100 | 165kb duplication | - | *hpx – sdhA* | - | | |

# MT345

| Replicate | Position | Frequency at passage 0 | Frequency at passage 4 | Type | Mutation | Gene | Product | Frequency in Global Dataset | Found in planktonically passaged populations |
|---|---|---|---|---|---|---|---|---|---|
| A | 3305000 | ~50 | ~75 | 770kb duplication | - | *lppX – Rv3639c* | - | | |
| B | 3305000 | ~50 | ~75 | 770kb duplication | - | *lppX – Rv3639c* | - | | |
| C | 3305000 | ~50 | ~75 | 770kb duplication | - | *lppX – Rv3639c* | - | | |

# MT72

| Replicate | Position | Frequency at passage 0 | Frequency at passage 4 | Type | Mutation | Gene | Product | Frequency in Global Dataset | Found in planktonically passaged populations |
|---|---|---|---|---|---|---|---|---|---|
| B | 3360339 | 0 | 84 | Non-synonymous | Q83R (CAG-CGG) | *ilvC* | KETOL-acid reductoisomerase | 0 | |
| C | 784878 | 9 | 39 | Non-synonymous | H20D (CAC-GAC) | *tuf* | elongation factor Tu | 0 | Yes |
| C | 2399030 | 0 | 32 | Non-synonymous | L104P (CTG-CCG) | *pyrD* | dihydroorotate dehydrogenase | 0 | |

# MT49

| Replicate | Position | Frequency at passage 0 | Frequency at passage 4 | Type | Mutation | Gene | Product | Frequency in Global Dataset | Found in planktonically passaged populations |
|---|---|---|---|---|---|---|---|---|---|
| A | 739751 | 0 | 86 | Non-synonymous | Y146C (TAT-TGT) | *mmaA1* | methoxy mycolic acid synthase | 0 | |
| A | 3336587 | 73 | 45 | Intergenic | +82/+209 (T-C) | *Rv2980 / ddlA* | conserved secreted protein / D-alanine ligase | N/A | |
| C | 3336587 | 73 | 39 | Intergenic | +82/+209 (T-C) | *Rv2980 / ddlA* | conserved secreted protein / D-alanine ligase | N/A | |

# MT540

| Replicate | Position | Frequency at passage 0 | Frequency at passage 4 | Type | Mutation | Gene | Product | Frequency in Global Dataset | Found in planktonically passaged populations |
|---|---|---|---|---|---|---|---|---|---|
| A | 524164 | 16 | 68 | Non-synonymous | G124S (GGC-AGC) | *Rv0435c* | ATPase | 0.015% | Yes |
| A | 782876 | 11 | 49 | Non-synonymous | V131G (GTC-GGC) | *fusA1* | elongation factor G | 0 | |
| A | 851966 | 0 | 33 | Non-synonymous | T120K (ACA-AAA) | *phoP* | two component system response transcriptional regulator | 0 | |
| A | 1292786 | 9 | 45 | Intergenic | +45/-12 (A-G) | *narH / narJ* | nitrate reductase beta chain / nitrate reductase delta chain | 0 | Yes |

| | | | | | | | | |
|---|---|---|---|---|---|---|---|---|
| B | 104838 | 17 | 72 | Non-synonymous | E126D (GAA-GAC) | *Rv0095c* | hypothetical protein | 0.60% |
| B | 1986846 | 29 | 88 | Intergenic | -176/+8 (C-T) | *Rv1754c / plcD* | hypothetical protein / phospholipase C | 0 |
| B | 3690947 | 0 | 90 | Intergenic | -9/-194 (A-G) | *Rv3303c / Rv3304* | NAD(P)H quinone reductase / hypothetical protein | 0.15% |
| C | 104838 | 17 | 80 | Non-synonymous | E126D (GAA-GAC) | *Rv0095c* | hypothetical protein | 0 |
| C | 1986846 | 29 | 100 | Intergenic | -176/+8 (C-T) | *Rv1754c / plcD* | hypothetical protein / phospholipase C | 0 |
| C | 3689529 | 0 | 98 | Synonymous | L470L (CTG-CTA) | *lpdA* | NAD(P)H quinone reductase | 0 |

***M. tb* loci with allele frequency changes >=30% under selection for biofilm growth.** Results are broken down by strain. The frequency of each SNP in a natural population sample is shown (Global Dataset, see Materials and Methods). Note: N/A in MT49 table indicates that a reliable natural population frequency could not be calculated for that variant. If an identical variant was also found in planktonically passaged populations of *M. tb*, this is indicated.

**Figure S3:**

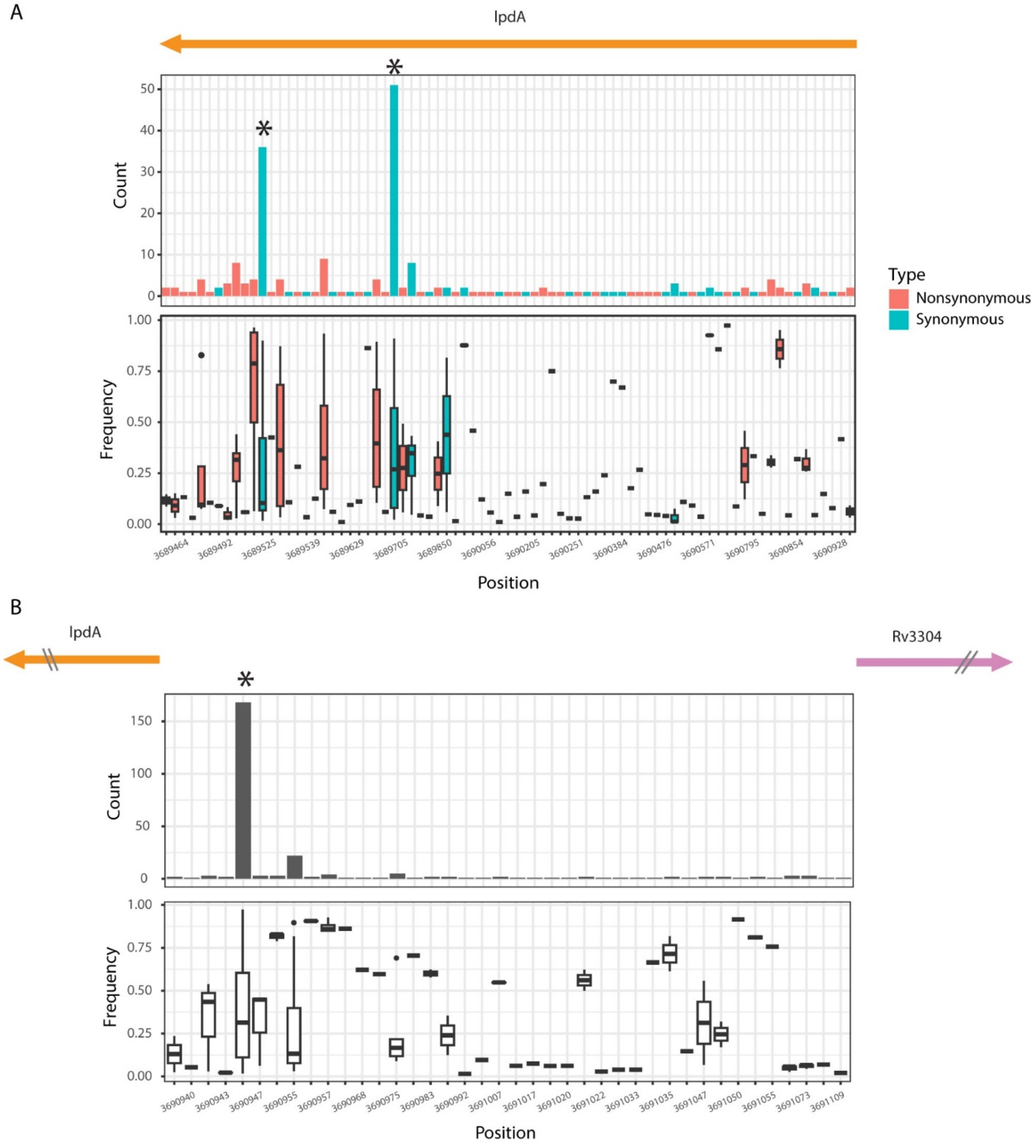

**Biofilm mutations under selection within hosts with TB.** Mutation counts (top) and within-host frequencies (bottom) of mutations in *lpdA* (A) and the intergenic region between *lpdA* and *Rv3304* (B) from Liu et al (2022)'s analysis of >50,000 samples. Asterisks indicate the mutations that emerged in our study during biofilm selection, including two synonymous variants from

MT31 and MT540 and an intergenic variant that arose independently in MT49 and MT540 in different rounds of the experiment.

**Table S3:**

| Experiment | Strain | Replicate | Position | Passage | | | | | | | Gene | Frequency in Global Dataset |
|---|---|---|---|---|---|---|---|---|---|---|---|---|
| | | | | **4B-4C** | **0** | **1** | **2** | **3** | **4** | **5** | | |
| **Competition** | 31-4B4C | A | 2447977 | 39 | 14 | 0 | 0 | 0 | 0 | 0 | *Rv2186c* | 0.0025% |
| | 31-4B4C | A | 3657010 | 0 | 0 | 19 | 84 | 100 | 100 | 100 | *fadE25* | 0 |
| | 31-4B4C | A | 3689763 | 22 | 36 | 60 | 93 | 100 | 100 | 100 | *lpdA* | 0.058% |
| | 31-4B4C | A | – | 56 | 39 | 24 | 6 | 3 | 3 | 2 | MMMC | NA |
| | 31-4B4C | B | 2447977 | 39 | 10 | 43 | 38 | 73 | 77 | 88 | *Rv2186c* | 0.0025% |
| | 31-4B4C | B | 3689763 | 22 | 48 | 17 | 12 | 0 | 0 | 0 | *lpdA* | 0.058% |
| | 31-4B4C | B | – | 56 | 37 | 63 | 67 | 89 | 87 | 90 | MMMC | NA |
| | 31-4B4C | C | 2447977 | 39 | 6 | 0 | 0 | 0 | 0 | 0 | *Rv2186c* | 0.0025% |
| | 31-4B4C | C | 3689763 | 22 | 48 | 81 | 86 | 100 | 100 | 94 | *lpdA* | 0.058% |
| | 31-4B4C | C | – | 56 | 31 | 12 | 8 | 3 | 2 | 5 | MMMC | NA |
| | **Strain** | **Replicate** | **Position** | **Passage** | | | | | | | **Gene** | **Frequency in Global Dataset** |
| | | | | **4** | **4.0** | **4.1** | **4.2** | **4.3** | **4.4** | **4.5** | | |
| **Extended Passaging** | 31-4B | A | 2447977 | 0 | 0 | 0 | 0 | 0 | 0 | 0 | *Rv2186c* | 0.0025% |
| | 31-4B | A | 3689763 | 53 | 51 | 69 | 100 | 92 | 88 | 62 | *lpdA* | 0.058% |
| | 31-4B | A | – | 25 | 30 | 17 | 4 | 4 | 8 | 27 | MMMC | NA |
| | 31-4B | B | 2447977 | 0 | 0 | 0 | 0 | 0 | 0 | 0 | *Rv2186c* | 0.0025% |
| | 31-4B | B | 3657010 | 0 | 5 | 0 | 49 | 70 | 66 | 97 | *fadE25* | 0 |
| | 31-4B | B | 3689763 | 53 | 52 | 68 | 88 | 96 | 96 | 98 | *lpdA* | 0.058% |
| | 31-4B | B | – | 25 | 28 | 20 | 6 | 2 | 6 | 2 | MMMC | NA |
| | 31-4B | C | 2447977 | 0 | 0 | 0 | 0 | 0 | 0 | 0 | *Rv2186c* | 0.0025% |
| | 31-4B | C | 3689763 | 53 | 54 | 55 | 74 | 71 | 74 | 97 | *lpdA* | 0.058% |
| | 31-4B | C | – | 25 | 27 | 22 | 17 | 15 | 14 | 4 | MMMC | NA |

***M. tb* loci with allele frequency changes >=30% following extended passaging (MT31-4B) or competition (MT31-4B4C).** The frequency of each SNP in a natural population sample is shown (Global Dataset, see Materials and Methods, SI Appendix 2).
