## Supplementary Data 2 for "Contingencies in biofilm adaptation of *Mycobacterium tuberculosis*"

Materials and Methods

**Storage and resuscitation of ancestral populations**

Between the first and second round of these experiments, the ancestral populations from round 1 were thawed and grown in planktonic cultures to create more freezer stocks. Briefly, freezer stocks were thawed and inoculated into Middlebrook 7H9 broth (HiMedia) containing 0.2% w/v glycerol, 10% v/v OADC supplement (oleic acid, albumin, D-glucose and catalase; Becton Dickinson), and 0.05% w/v Tween-80. Planktonic cultures were incubated at 37C with shaking until grown to an OD600 ~1. These planktonic cultures were then frozen into cryovials which would be used as the ancestors for round 2. Ancestral populations were resuscitated from freezer stocks and grown as pellicles as previously described (Smith et al. 2022). Pellicles were grown by inoculating 250 uL of planktonic culture (grown as described above) into 25 mL Sauton's medium (for 1 L: 0.5 g $KH_2PO_4$, 0.5 g $MgSO_4$, 4 g L-asparagine, 2 g citric acid, 50 mg ammonium iron (III) citrate, 60 mL glycerol, adjust pH to 7.0 with NaOH) containing 0.1% w/v $ZnSO_4$ and incubated at 37°C with 5% $CO_2$ supplementation, without shaking. Biofilms were grown as previously described (Kulka, Hatfull, and Ojha 2012): 250 mL bottles (Corning, 430281) were incubated with a tight-fitting cap for 3 weeks and then with a loose-fitting cap for 2 additional weeks for a total of 5 weeks of growth.

**Pellicle passaging and phenotyping**

Ancestral populations were split into three independently evolving replicates per strain, and passaged four times per replicate as previously described (Smith et al. 2022). Briefly, 0.3 g (wet weight) of mature pellicle was inoculated into a fresh bottle containing 25 mL Sauton's and grown as described above. At each passage point pellicle growth was documented with photos, through categorical phenotyping (Figure S1) and with wet weight measurements. Wet weight measurements were taken by removing spent media beneath the pellicle, weighing and subtracting the tare weight of the empty bottle.

**Extended passaging and competition of MMMC duplication and *lpdA***

Frozen stocks of the fourth passage of MT31 replicates B and C were thawed and grown planktonically as described above. For the extended passaging experiment MT31-4B was grown as a pellicle (Passage 0) and passaged as described above. For the competition experiment, equal volumes of MT31-4B and MT31-4C were inoculated into Sauton's for pellicle growth (Passage 0) and passaged as described above.

**DNA extraction and sequencing**

At each passage point genomic DNA (gDNA) was extracted from the whole pellicle for pooled sequencing. gDNA was extracted using one of two methods: a modified Qiagen DNeasy Blood and Tissue kit protocol which we previously described (Smith et al. 2022) or a phenol-chloroform extraction. For phenol-chloroform extractions, pellicles were transferred to conical tubes before being spun at 5000 x *g* for 10 min and the resulting supernatant discarded. The pellet was resuspended in 1mL of PBS and 450 uL of the suspension was transferred to a screw-cap tube with 50 uL of lysozyme (100 mg/mL) and incubated overnight at 37°C. Next 10 uL of RNase (10 mg/mL) was added and samples were incubated at room temperature for 30 min. 70 uL of 10% SDS and 10 uL of Proteinase K were added before incubating at 65°C for 20 min. 100 uL of 5 M NaCL was added and samples were vortexed before adding 100 uL of NaCl-CTAB (4.1 g NaCl in 80 mL water, add 10 g N-cetyl-N,N,N,-trimethyl ammonium bromide [CTAB], adjust volume to 100 mL with distilled water, warmed in 65°C water bath prior to use). Samples were vortexed and incubated at 65°C for 10 min. Next, 750 uL of chloroform/iso-amyl alcohol (24:1) was added before vortexing, and spinning in a tabletop centrifuge at maximum speed for 10 min. After centrifugation the aqueous phase (top layer) was carefully removed and added to a new tube along with 750 uL of isopropanol. Samples were gently inverted before being placed in a -20°C freezer overnight. Next, samples were spun in a tabletop centrifuge at maximum speed for 20 min and supernatant was discarded before washing with 1 mL of cold 70% ethanol. Following a 5 min spin the supernatant was discarded and pellets were left to dry at room temp for 30 min. Finally, the gDNA pellet was gently dissolved (without pipetting) in RNase free water. gDNA samples were sent to the University of Wisconsin-Madison Biotechnology Center or to SeqCoast for library preparation and sequencing using paired-end 150 bp reads.

**SEM**

For SEM experiments, *M. tb* strains were grown as pellicle biofilms for 5-7 weeks before a piece of pellicle was placed on poly-L-lysine-treated plastic coverslips (13 mm, Thermonax plastic for cell culture). These pellicle pieces were then fixed overnight in a solution of 4% formaldehyde, 2% glutaraldehyde in Dulbecco's phosphate-buffered saline (-calcium, -magnesium) (DPBS) (Hyclone Laboratories Inc, Logan, UT). Following overnight fixation, samples were treated with DPBS. Next, samples were treated with 1% osmium tetroxide for 1 hr. After treatment with osmium tetroxide, samples were again washed with DPBS before undergoing sequential ethanol dehydration. Following ethanol dehydration, samples underwent critical point drying. Finally, samples were placed on aluminum stubs and sputter-coated with 20 nm platinum. The samples were imaged at 3 kV by a Zeiss GeminiSEM 450 SEM.

**Reference guided assembly and variant calling**

Genome assemblies were performed as previously described (Smith et al. 2022). Briefly, an in-house reference-guided mapping pipeline was used (https://github.com/pepperell-lab/RGAPepPipe_MAY) to produce pooled-sequencing alignments. We used Popoolation2 v1.201 (Kofler, Pandey, and Schlötterer 2011) along with in-house scripts (https://github.com/pepperell-lab/mtb_ExpEvo) to identify mutations that changed significantly in frequency (>= 30%) over the course of our experiment. Variants were filtered for quality by a minimum coverage of 20x, a minimum alternate allele count of 5 and a minimum allele frequency of 5%. We removed variants in repetitive gene families (PE, PPE, and PE-PGRS).

**Structural variant analysis**

Duplications in MMMC family were identified using sliding window coverage plots as previously described (Smith et al. 2022). The bounds of each duplication were estimated by identifying windows with >1.25x relative coverage. The frequency of duplications within each population was estimated by averaging relative coverage within duplication bounds and subtracting 1 (ie an average relative coverage of 1.5x corresponds to a duplication frequency of ~50%).

**Frequency in global dataset**

We estimated the frequency of mutations identified in this study in a global dataset of *M. tb* genomes based on a previously described method (Smith et al. 2022). Using a searchable compact bit-sliced signature (COBS) index of bacterial genomes curated from the European Nucleotide Archive (Blackwell et al. 2022). We used the Python interface to search the COBS index (Bingmann et al. 2019), using scripts available at https://github.com/pepperell-lab/Mtb_COBS. Our query sequences included 50bp in either direction of the SNP of interest, with a k-mer matching threshold of 1. The frequency of isolates with each SNP in the database was calculated by dividing the number of isolates with the SNP by the number of *M. tb* genomes in the dataset, which we previously determined to be ~40k (Smith et al. 2022).

A dataset of all unfixed mutations identified in their analysis of >50,000 *M. tb* isolates (Liu et al, 2022) was provided by the authors of the study. SNPs within *lpdA* and in the intergenic region between *lpdA* and *Rv3304* were isolated, and the number of mutations at each position was plotted along with the frequency of the mutation in the source population.

**Transformants**

The *lpdA* and *glpD2* genes were overexpressed in *M. tb* at the L5 site using the integrative, constitutive overexpression plasmid CT94 from the Sarah Fortune lab. The *lpdA* gene was amplified and modified to include NdeI and HindIII digestion sites using forward primer CCGCATGCTTAATTAAGAAGGAGATATACA-CGCCGAGCTAGGTTATGGGCT and reverse primer GACCTCTAGGGTCCCCAATTAATTAGCTAA-CCTGGATTGGGTTGCTCACGA. The *glpD2* gene was amplified and modified to NdeI and HindIII digestion sites using forward primer CCGCATGCTTAATTAAGAAGGAGATATACA-TAACTGACAGGAGCCGGTTTC and reverse primer GACCTCTAGGGTCCCCAATTAATTAGCTAA-CGACATACCCGCTTGGCGTAG. After amplification with Platinum SuperFi Green PCR Master Mix (Invitrogen, Waltham, MA), the *lpdA* and *glpD2* PCR products were digested with NdeI and HindIII (NEB, Ipswich, MA). The digested products were then ligated with a NdeI and HindIII cut CT94 plasmid using Gibson assembly (NEB, Ipswich, MA). The assembly products containing *lpdA* or *glpD2* and the CT94 plasmid were then heat-shock transformed into DH5α E. coli cells and selected for on LB plates with 50 ug/ml kanamycin overnight. Plasmid integration was verified via NdeI and HindIII digestion. The *lpdA* and *glpD2* overexpression plasmids were then electroporated into electrocompetent *M. tb* strain H37Rv along with the ancestral clinical strains, with the CT94 plasmid electroporated as a control. Transformants were recovered in 2 ml of fresh 7H9 for 24-48 h at 37°C, and then struck on 7H10 plates containing 25 ug/ml kanamycin.
